# Spoink, a LTR retrotransposon, invaded *D. melanogaster* populations in the 1990s

**DOI:** 10.1101/2023.10.30.564725

**Authors:** Riccardo Pianezza, Almorò Scarpa, Prakash Narayanan, Sarah Signor, Robert Kofler

## Abstract

During the last few centuries *D. melanogaster* populations were invaded by several transposable elements, the most recent of which was thought to be the *P*-element between 1950 and 1980. Here we describe a novel TE, which we named *Spoink*, that has invaded *D. melanogaster*. It is a 5216nt LTR retrotransposon of the Ty3/gypsy superfamily. Relying on strains sampled at different times during the last century we show that *Spoink* invaded worldwide *D. melanogaster* populations after the *P*-element between 1983 and 1993. This invasion was likely triggered by a horizontal transfer from the *D. willistoni* group, much as the *P*-element. *Spoink* is probably silenced by the piRNA pathway in natural populations and about 1/3 of the examined strains have an insertion into a canonical piRNA cluster such as *42AB*. Given the degree of genetic investigation of *D. melanogaster* it is surprising that *Spoink* was able to invade unnoticed.

## Introduction

Transposable elements (TEs) are short genetic elements that can increase in copy number within the host genome. They are abundant in most organisms and can make up the majority of some genomes, i.e. maize where TEs constitute 83% of the genome [Schnable et al., 2009]. There are two classes of TEs which transpose by different mechanisms - DNA transposons which replicate by directly moving to a new genomic location in a ‘cut and paste’ method, and retrotransposons which replicate through an RNA intermediate in a ‘copy and paste’ method [Kapitonov and Jurka, 2003, Finnegan, 1989, Wicker et al., 2007]. From humans to flies, more genetic variation (in bp) is due to repetitive sequences such as transposable elements than all single nucleotide variants combined [Chakraborty et al., 2021]. Although some TEs, such as *R1* and *R2* elements, may benefit hosts [Eickbush and Eickbush, 1995, Nelson et al., 2023] most TE insertions are thought to be deleterious [Elena et al., 1998, Pasyukova et al., 2004]. Host genomes have therefore evolved an elaborate system of suppression frequently involving small RNAs [Sarkies et al., 2015]. Suppression of TEs in *Drosophila* relies upon small RNAs termed piRNAs are cognate to TE sequences [Brennecke et al., 2007, Gunawardane et al., 2007, Brennecke et al., 2008]. These small RNAs bind to PIWI clade proteins and mediate the degradation of TE transcripts and the formation of heterochromatin silencing the TE [Brennecke et al., 2007, Le Thomas et al., 2013, 2014a,b, Yamanaka et al., 2014, Andreev et al., 2022, Rangan et al., 2011]. However, while host defenses quickly adapt to new transposon invasions, TEs can escape silencing through horizontal transfer to new, defenseless, genomes [Peccoud et al., 2017, Scarpa et al., 2023, Kofler et al., 2015, Signor et al., 2023].

This horizontal transfer allows TEs to colonize the genomes of novel species [Zhang et al., 2020, Zanni et al., 2013, Signor et al., 2023, Schaack et al., 2010, Peccoud et al., 2017]. The first well-documented instance of horizontal transfer of a TE was the *P*-element, which spread from *D. willistoni* to *D. melanogaster* [Daniels et al., 1990]. Following this horizontal transfer the *P*-element invaded natural *D. melanogaster* populations between 1950 and 1980 [Anxolabéhère et al., 1988, Kidwell, 1983]. It was further realized that the *I*-element, *Hobo* and *Tirant* spread in *D. melanogaster* populations earlier than the P-element, between 1930 and 1960 [Kidwell, 1983, Periquet et al., 1989, Schwarz et al., 2021]. The genomes from historical *D. melanogaster* specimens collected about two hundred years ago, recently revealed that *Opus, Blood*, and *412* spread in *D. melanogaster* populations between 1850 and 1933 [Scarpa et al., 2023]. In total, it was suggested that seven TEs invaded *D. melanogaster* populations during the last two hundred years where one invasion (the *P*-element) was triggered by horizontal transfer from a species of the *willistoni* group and six invasions by horizontal transfer from the *simulans* complex [Schwarz et al., 2021, Scarpa et al., 2023, Loreto et al., 2008, Daniels et al., 1990, Simmons, 1992, Blumenstiel, 2019].

It was, however, widely assumed until now that the *P*-element invasion, which occurred between 1950-1980, was the last and most recent TE invasion in *D. melanogaster* [Kidwell, 1983, Anxolabéhère et al,. 1985, Bonnivard and Higuet, 1999, Schwarz et al,. 2021, Scarpa et al,. 2023]. Here we report the discovery of Spoink, a novel TE which invaded worldwide *D. melanogaster* populations between 1983 and 1993, i.e. after the invasion of the *P*-element. *Spoink* is a LTR retrotransposon of the Ty3/gypsy group. We suggest that the *Spoink* invasion in *D. melanogaster* was triggered by horizontal transfer from a species of the *willistoni* group, similarly to the *P*-element invasion in *D. melanogaster*. In a model species as heavily investigated as *D. melanogaster* it is surprising that *Spoink* was able to invade undetected.

## Results

Previous work showed that at least seven TE families invaded *D. melanogaster* populations during the last two hundred years [Scarpa et al., 2023, Schwarz et al., 2021, Kidwell, 1983]. To explore whether additional, hitherto poorly characterised TEs could have invaded *D. melanogaster*, we investigated long-read assemblies of recently collected *D. melanogaster* strains [Rech et al., 2022] using a newly assembled repeat library [Chakraborty et al., 2021]. Interestingly we found differences in the abundance of “gypsy 7 DEl” between the reference strain *Iso-1* and more recently collected *D. melanogaster* strains (Supplementary table S1). To better characterize this TE, we generated a consensus sequence based on the novel insertions and checked if this consensus sequence matches any of the repeats described in repeat libraries generated for *D. melanogaster* and related species [Quesneville et al., 2005, Rech et al., 2022, Chakraborty et al., 2021, Srivastav et al., 2023, Ellison and Cao, 2020]. A fragmented copy of this TE, with just one of the two LTRs being present, was reported by Rech et al. [2022] (0.13% divergence; “con41 UnFmcl001 RLX-incomp”; Supplementary table S2). The next best hits were *gypsy7 Del, gypsy2 DSim, micropia* and *Invader6* (18-30% divergence; Supplementary table S2). Given this high sequence divergence from previously described TE families and the fact that this novel TE belongs to an entirely different superfamily/group than *gypsy7* (see below), we decided to give this TE a new name. We call this novel TE *”Spoink”* inspired by a Pokémon that needs to continue jumping to stay alive.

*Spoink* is an LTR retrotransposon with a length of 5216 bp and LTRs of 349 bp (fig. 1A). At positions 4639-4700 *Spoink* contains a poly-A tract with a few mismatches. *Spoink* encodes a 695 aa putative *gag-pol* polyprotein. Ordered from the N-to the C-terminus the conserved domains of the polyprotein are zinc knuckle (likely part of *gag* ; e-value *e* = 4.5*e* − 03), retropepsin (*e* = 1.3*e* − 05), reverse transcriptase of LTR (*e* = 1.8*e* − 59), RNase HI of Ty3/gypsy elements (*e* = 2.9*e* − 38), integrase zinc binding domain (*e* = 2.9*e* − 16) and integrase core domain (*e* = 4.5*e* − 10). *Spoink* lacks an *env* protein. The order of these domains, with the integrase downstream of the reverse transcriptase, is typical for Ty3/gypsy transposons [Eickbush and Malik, 2002].

**Figure 1.**
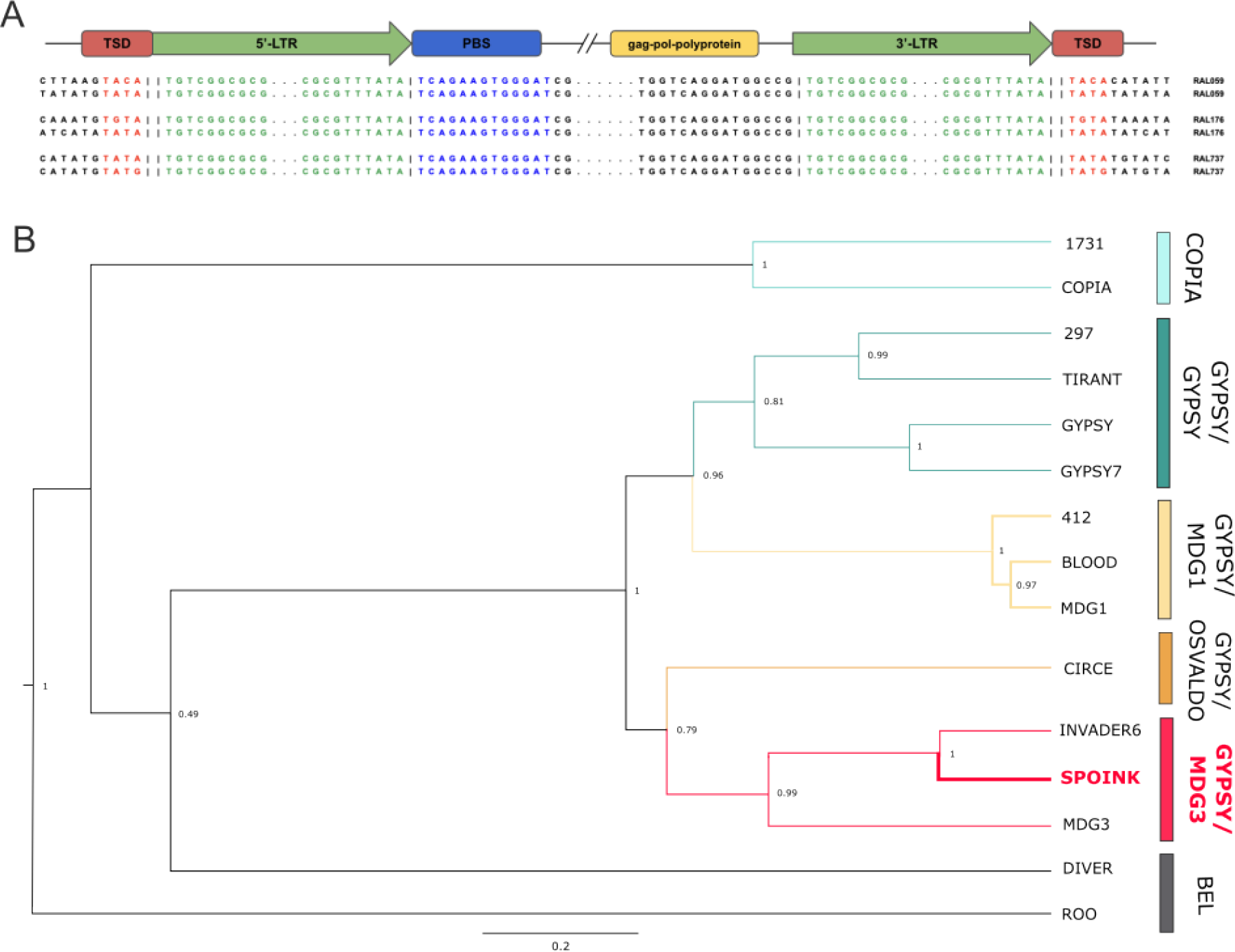
*Spoink* is a novel TE of the Ty3/gypsy superfamily. A) Overview of the composition of *Spoink*. Features are shown in color and the alignments show the sequences around the main features of *Spoink* for two insertions in each of three different long-read assemblies of *D. melanogaster*. B) Phylogenetic tree based on the reverse-transcriptase domain of *pol* for *Spoink* and several other LTR transposons. Multiple families have been picked for each of the main superfamilies/groups of LTR transposons [Kapitonov and Jurka, 2003]. Our data suggest that *Spoink* is a member of the gypsy/mdg3 group.

A phylogeny based on the reverse transcriptase domain of different TE families suggests that *Spoink* is a member of the gypsy/mdg3 superfamily/group of LTR retrotransposons (fig. 1B; [Kapitonov and Jurka, 2003]). As expected for members of the Ty3/gypsy superfamily *Spoink* generates a target site duplication of 4 bp and it has an insertion motif enriched for ATAT (fig. 1A; [Kapitonov and Jurka, 2003, Linheiro and Bergman, 2012]). However, *Spoink* differs from what is expected for the Ty3/gypsy superfamily in several ways. First, the encoded *gag-pol* polyprotein is atypical for Ty3/gypsy transposons [Eickbush and Malik, 2002]. Second, the predicted primer binding site of *Spoink* directly follows the LTR, whereas typically for Ty3/gypsy there is a shift of 5-8nt (fig. 1A; [Kapitonov and Jurka, 2003]). Third, the LTR motif is TG…TA which is different from the TG…CA motif usually reported for *gypsy* TEs [Kapitonov and Jurka, 2003].

Finally we investigated the genomic distribution of *Spoink* insertions in long-read assemblies of *D. melanogaster* strains collected ≥ 2003 [Rech et al., 2022]. In total, these assemblies contains 481 full-length (*>* 80% length with at least one LTR) insertions of *Spoink* (on the average 16 per genome). Unlike the *P*-element which has a strong insertion bias into promoters, *Spoink* insertions are mostly found in introns and intergenic regions (Supplementary fig. S1). 54% of the *Spoink* insertions are in 201 different genes. Interestingly we found 7 independent *Spoink* insertions in *Myo83F*.

To summarize we characterized a novel LTR-retrotransposon of the Ty3/gypsy superfamily in the genome of *D. melanogaster* that we call *Spoink*.

### *Spoink* recently invaded worldwide *D. melanogaster* populations

To substantiate our hypothesis that *Spoink* recently invaded *D. melanogaster* we used three independent approaches: Illumina short read data, long-read assemblies, and PCR/Sanger sequencing. First we aligned short reads from a strain collected in 1958 (*Hikone-R*) and a strain collected in 2015 (*Ten-15*) [Rech et al., 2022, Schwarz et al., 2021] to the consensus sequence of *Spoink* using DeviaTE [Weilguny and Kofler, 2019]. DeviaTE estimates the abundance of *Spoink* insertions by normalizing the coverage of *Spoink* to the coverage of single-copy genes. Furthermore, DeviaTE is useful for generating an intuitive visualization of the abundance and composition (i.e. SNPs, indels, truncations) of *Spoink* in samples. We found that only a few degraded reads aligned to *Spoink* in the 1950’s strain (*Hikone-R*) whereas many reads covered the sequence of *Spoink* in the more recently collected strain *Ten-15* (fig. 2A). There were also very few SNPs or indels in the recently collected strain suggesting that most insertions have a very similar sequence (fig. 2A). This observation holds true when multiple old and young *D. melanogaster* strains are analysed (Supplementary fig. S1).

**Figure 2.**
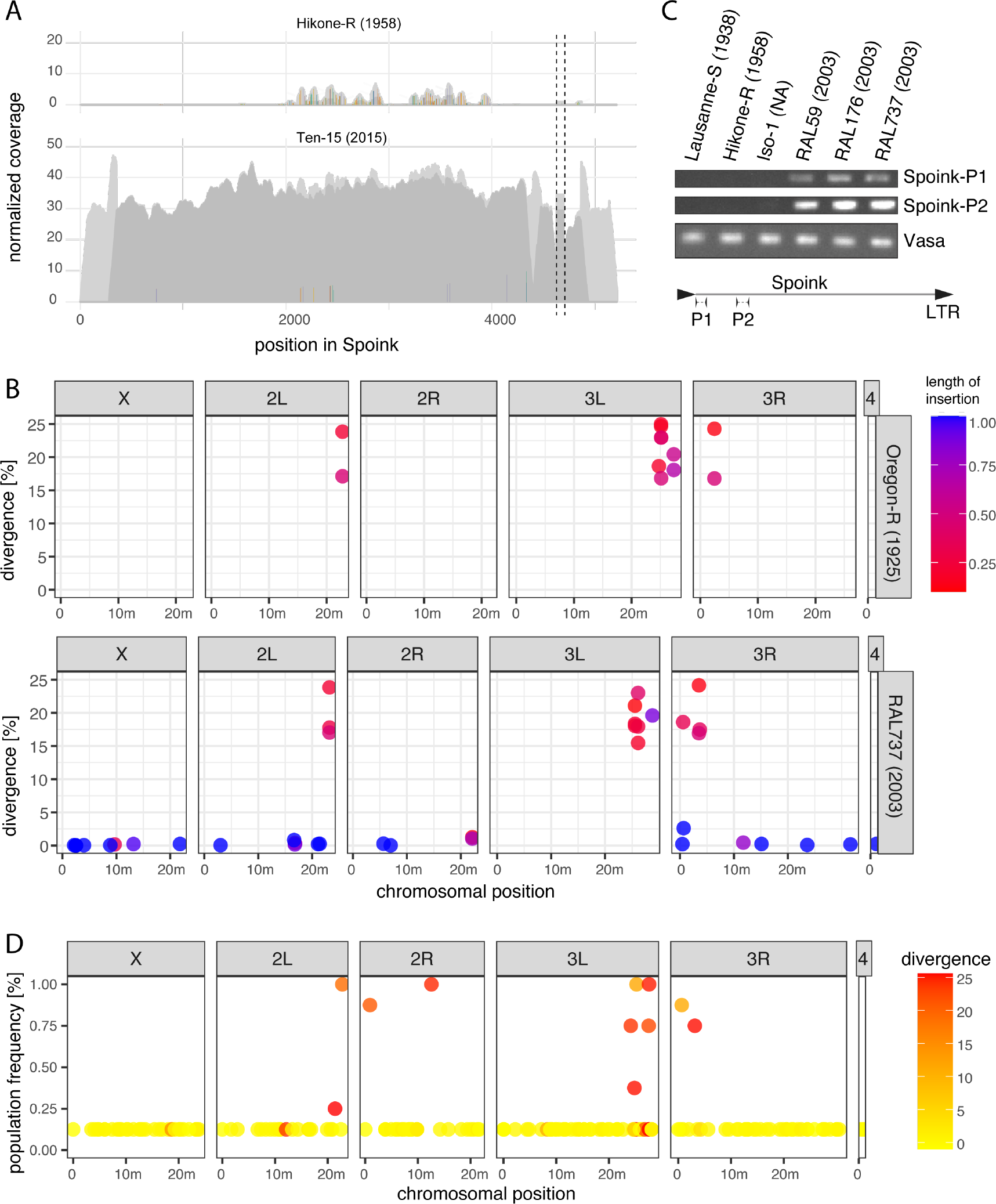
*Spoink* invaded *D. melanogaster*. A) DeviaTE plots of *Spoink* for a strain collected in 1954 (*Hikone-R*) and a strain collected in 2015 (*Ten-15*). Short reads were aligned to the consensus sequence of *Spoink* and the coverage was normalized to the coverage of single-copy genes. The coverage based on uniquely mapped reads is shown in dark grey and light grey is used for ambiguously mapped reads. Single-nucleotide polymorphisms (SNPs) and small internal deletions (indels) are shown as colored lines. The coverage was manually curbed at the poly-A track (between dashed lines). B) Insertions with a similarity to the consensus sequence of *Spoink* in the long-read assemblies of *Oregon-R* (collected around 1925) and the more recently collected strain *RAL737* (2003). C) PCR results for two *Spoink* primer pairs (for location of primers see sketch at bottom) and one primer pair for the gene *vasa. Spoink* is absent in old strains (*Lausanne-S, Hikone-R* and *iso-1*) and present in more recently collected strains (*RAL59, RAL176, RAL737*). D) Population frequency of *Spoink* insertions in long-read assemblies of strains collected in 2003 from Raleigh [Rech et al., 2022]. Note that highly diverged insertions are largely segregating at a high frequency while canonical *Spoink* insertions mostly segregate at a low frequency.

Next we investigated the abundance of *Spoink* in long-read assemblies of a strain collected in 1925 (*Oregon-R*) and a strain collected in 2003 (*RAL737*). We found solely highly diverged and fragmented copies of sequences with similarity to *Spoink* in *Oregon-R* (fig. 2B). These degraded fragments were mostly found near the centromeres of *Oregon-R*. Investigating the identity of these degraded fragments of *Spoink* in more detail we found that they largely match with short and highly diverged fragments of *Invader6, micropia* and the *Max-element* (Supplementary table S3). In addition to these degraded fragments, the more recently collected strain *RAL737* also carries a large number of full-length insertions with a high similarity to the consensus sequence of *Spoink* (henceforth canonical *Spoink* insertions; fig. 2B). The canonical *Spoink* insertions are distributed all over the chromosomes of *RAL737* (fig. 2B). This observation is again consistent when several long-read assemblies of old and young *D. melanogaster* strains are analysed (Supplementary fig. S2).

Finally we used PCR to test whether *Spoink* recently spread in *D. melanogaster*. We designed two PCR primer pairs for *Spoink* and, as a control, one primer pair for *vasa* (fig. 2C; bottom panel). The *Spoink* primers amplified a clear band in three strains collected 2003 in Raleigh but no band was found in earlier collected strains, including the reference strain of *D. melanogaster, Iso-1* (fig. 2C). We sequenced the fragments amplified by the *Spoink* primers using Sanger sequencing and found that the sequence of the six amplicons matches with the consensus sequence of *Spoink* (Supplementary fig. S3).

Finally we investigated the population frequency of canonical and degraded *Spoink* insertions. Using the long-read assemblies of eight strains collected in 2003 in Raleigh we computed the population frequency of different *Spoink* insertions. We found that canonical *Spoink* insertions (*<* 5% divergence) are largely segregating at a low population frequency, as expected for recently active TEs (fig. 2D). While several degraded fragments that were annotated as *Spoink* are private, there were many at a higher population frequency as expected for older sequences (fig. 2D).

In summary our data suggest that *Spoink* recently spread in *D. melanogaster* and that degraded fragments with some similarity to *Spoink* are present in all investigated *D. melanogaster* strains. These degraded fragments may be the remnants of more ancient invasions of TEs sharing some sequence similarity with *Spoink*.

### Timing the *Spoink* invasion

Next we sought to provide a more accurate estimate of the time when *Spoink* spread in *D. melanogaster*. First we generated a rough timeline of the *Spoink* invasion using *D. melanogaster* strains sampled during the last two hundred years. We estimated the abundance of *Spoink* in these strains using DeviaTE [Weilguny and Kofler, 2019]. As reference we also estimated the abundance of the *P*-element, which is widely assumed as to be the most recent TE that invaded *D. melanogaster* populations [Schwarz et al., 2021, Anxolabéhère et al., 1988]. *Spoink* was absent from all strains collected ≤ 1983 but present in strains collected ≥ 1993 (fig. 3A). By contrast our data suggest that the *P*-element was absent in the strains collected ≤ 1962 but present in strains collected ≥ 1967 (fig. 3A). This is consistent with previous works suggesting that the *P*-element invaded *D. melanogaster* between 1950 and 1980 [Kidwell, 1983, Anxolabéhère et al., 1985, Bonnivard and Higuet, 1999, Scarpa et al., 2023]. Our data thus suggest that *Spoink* invaded *D. melanogaster* after the *P*-element invasion. To investigate the timing of the invasion in more detail we estimated the abundance of *Spoink* in short-read data of 183 strains collected between 1960 and 2015 from different geographic regions using DeviaTE (Supplementary table S4; data from [Grenier et al., 2015, Schwarz et al., 2021, Long et al., 2013, Lange et al., 2021, Rech et al,. 2022]). The analysis of these 183 strains supports the view that Spoink was largely absent in strains collected ≤ 1983 but present in strains collected ≥ 1993 (fig. 3B). However there are two outliers. *Spoink* is present in one strain collected in 1979 in Providence (USA), which could be due to a contamination of the strain. On the other hand *Spoink* is absent in one strain collected ≥ 1993 in Zimbabwe (fig. 3B). As *Spoink* was present in six other strains collected in 1993 from Zimbabwe, it is feasible that *Spoink* was still spreading in populations from Zimbabwe around 1993. The strains supporting the absence of *Spoink* prior to 1983 were collected from Europe, America, Asia and Africa while the strains supporting the presence of *Spoink* after 1993 were collected from all five continents (Supplementary table S4).

**Figure 3.**
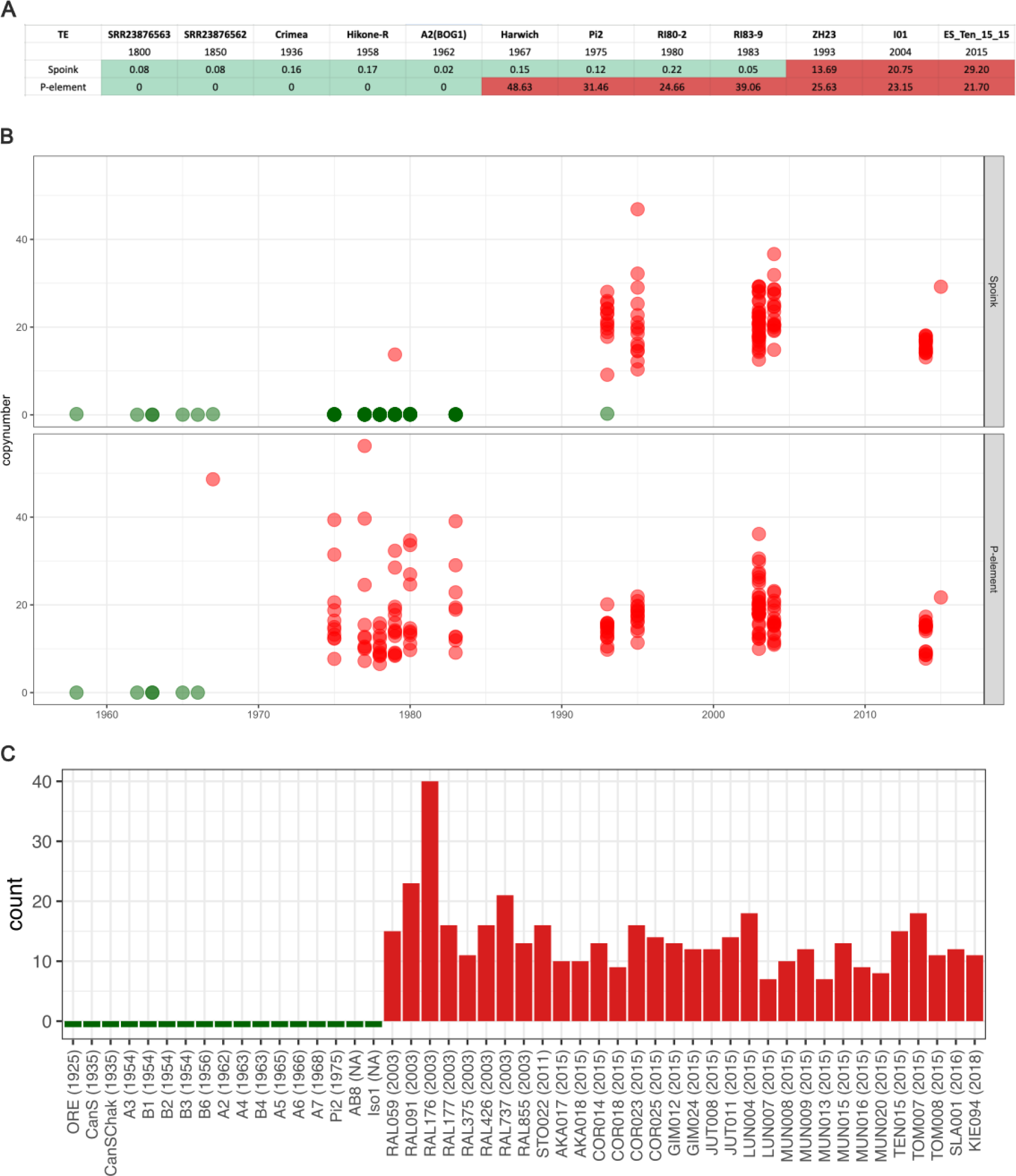
*Spoink* invaded *D. melanogaster* between 1983 and 1993 after the invasion of the *P*-element. A) Rough timeline of the *Spoink* and *P*-element invasion based on different strains sampled during the last two hundred years. B) Timeline of the *Spoink* and *P*-element invasion based on 183 strains sampled between 1960 and 2015 C) Abundance of canonical *Spoink* insertions (*>* 80% length and *<* 5% divergence) in long-read assemblies of *D. melanogaster* strains collected between 1925 and 2018.

Finally we estimated the abundance of *Spoink* in 49 long-read assemblies of strains collected during the last 100 years (Supplementary table S5; [Chakraborty et al., 2019, Wierzbicki et al., 2021, Hoskins et al., 2015, Rech et al., 2022]). We used RepeatMasker [Smit et al., 1996-2010] to estimate the abundance of canonical *Spoink* insertions (*>* 80% length and *<* 5% divergence) in these strains. Canonical *Spoink* insertions were absent in strains collected before 1975 but present in all long-read assemblies of strains collected after 2003 (fig. 3C). The strains of the assemblies supporting the absence of canonical *Spoink* insertions were collected from America, Europe, Asia, and Africa whereas the strains showing the presence of *Spoink* were largely collected from Europe, though genomes from North America and Africa are also represented (Supplementary table S5).

In summary we conclude that *Spoink* invaded worldwide populations of *D. melanogaster* approximately between 1983 and 1993. Moreover, the *Spoink* invasion is more recent than the *P*-element invasion.

### Geographic heterogeneity in the *Spoink* composition

Previous work showed that the composition of TEs within a species may differ among geographic regions [Schwarz et al., 2021, Scarpa et al., 2023]. Such geographic heterogeneity could result from founder effects occurring during the geographic spread of a TE. For example, a TE spreading in a species with a cosmopolitan distribution such as *D. melanogaster* may need to overcome geographic obstacles such as oceans and deserts. The few individuals that overcome these obstacles, thereby spreading the TE into hitherto naive populations, may carry slightly different variants of the TE than the source populations. These distinct variants will than spread in the new population. Such founder effects during the invasion may lead to a geographically heterogeneous composition of a TE within a species. For example, for the retrotransposon *Tirant*, individuals sampled from Tasmania carry distinct variants, while for *Blood* and *Opus* individuals from Zimbabwe are distinct from the other populations. To investigate whether we find such geographic heterogeneity we analysed the *Spoink* composition in the Global Diversity Lines (GDL), which comprise 85 *D. melanogaster* strains sampled after 1988 from five different continents (Africa - Zimbabwe, Asia - Beijing, Australia - Tasmania, Europe - Netherlands, America - Ithaca; [Grenier et al., 2015]). Except for a single strain from Zimbabwe all GDL strains harbour *Spoink* insertions (supplementary fig. S4). We estimated the allele frequency of SNPs in *Spoink*, where a SNP refers to a variant among dispersed copies of *Spoink*. The allele frequency estimate thus reflects the composition of *Spoink* within a particular strain. To summarize differences in the composition among the GDL strains we used UMAP [Diaz-Papkovich et al., 2021]. We found that the composition of *Spoink* varies among regions where three distinct groups can be distinguished: Tasmania, Bejing/Ithaca and Netherlands/Zimbabwe (supplementary fig. S4). It is interesting that clusters are formed by geographically distant populations such as Bejing (Asia) and Ithaca (America). We speculate that human activity, where flies might for example hitchhike with merchandise, could be responsible for this pattern. In summary, we found a geographically heterogeneous composition of *Spoink* which is likely due to founder effects occurring during the spread of this TE.

### *Spoink* is silenced by the piRNA pathway in natural populations

The host defence against TEs in *Drosophila* is based on small RNAs termed piRNAs. These piRNAs bind to PIWI clade proteins and silence a TE at the transcriptional as well as the post-transcriptional level [Brennecke et al., 2007, Gunawardane et al., 2007, Sienski et al., 2012, Le Thomas et al., 2013]. To test whether *Spoink* is silenced in *D. melanogaster* populations we investigated small RNA data from the GDL lines [Luo et al., 2020]. Small RNA were sequenced for 10 out of the 84 GDL lines such that two strains were picked from each of the five continents [Luo et al., 2020].

We find piRNAs mapping along the sequence of *Spoink* in the GDL strain *I17* which was collected in 2004 but not in the strain *Lausanne-S* which was sampled around 1938 (fig. 4A; [Lindsley and Grell, 1968]). piRNAs mapping to *Spoink* we further found for all 10 GDL strains (Supplementary fig. S5).

**Figure 4.**
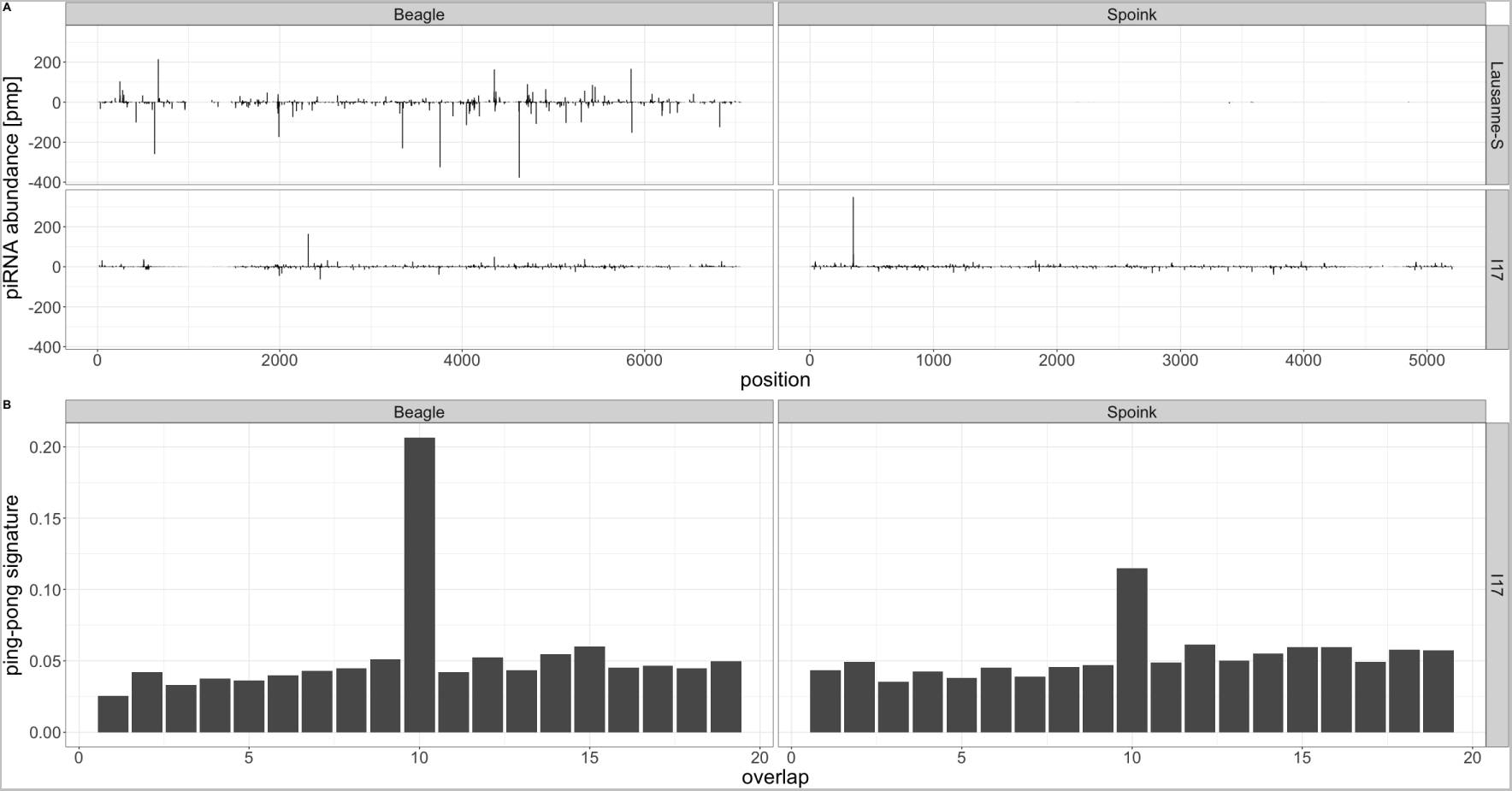
A piRNA based defence against *Spoink* emerged in *D. melanogaster* A) piRNAs mapping to *Spoink* in a strain sampled 1938 (*Lausanne-S*) and 2004 (*I17*). The transposon *Beagle* is included as reference. Solely the 5’ positions of piRNAs are shown and the piRNA abundance is normalized to one million piRNAs. Sense piRNAs are shown on the positive y-axis and antisense piRNAs on the negative y-axis. B) Ping-pong signature for the piRNAs mapping to *Spoink* and *Beagle* in the *D. melanogaster* strain *I17* (2004).

An important feature of germline piRNA activity in *D. melanogaster* is the ping-pong cycle [Brennecke et al., 2007, Gunawardane et al., 2007]. An active ping-pong cycle generates a characteristic overlap between the 5’ positions of sense and antisense piRNAs, i.e. the ping-pong signature. Computing a ping-pong signature thus requires several overlapping sense and antisense piRNAs. Since the amount of piRNAs was too low we could not compute a ping-pong signature for the strain *Lausanne-S* (collected in 1938; see above). However we found a pronounced ping-pong signature in all 10 GDL samples (fig. 4B; Supplementary fig. S5).

It is an important open question as to which events trigger the emergence of piRNA based host defence. The prevailing view, the trap model, holds that the piRNA based host defence is initiated by a copy of the TE jumping into a piRNA cluster [Bergman et al., 2006, Malone and Hannon, 2009, Zanni et al., 2013, Goriaux et al., 2014, Yamanaka et al., 2014]. If this is true we expect *Spoink* insertions in piRNA clusters in each of the long-read assemblies of the recently collected *D. melanogaster* strains [Rech et al., 2022]. We identified the position of piRNA clusters in these long-read assemblies based on unique sequences flanking the piRNA clusters [Wierzbicki et al., 2021]. Interestingly we found an extremely heterogeneous abundance of *Spoink* insertions in piRNA clusters, where some strains (e.g. *RAL176*) have up to 14 cluster insertions whereas 18 out of 31 strains did not have a single cluster insertion. Three of the cluster insertions were into *42AB*, which usually generates the most piRNAs [Brennecke et al., 2007, Srivastav et al., 2023]. It is an important open question whether such a heterogeneous distribution of *Spoink* insertions in piRNA clusters is compatible with the trap model [Kofler, 2019, Wierzbicki and Kofler, 2023]. In summary we found that *Spoink* is silenced by the piRNA pathway but the number of *Spoink* insertions in piRNA cluster is very heterogeneous among strains.

### Origin of *Spoink*

The invasion of *Spoink* in *D. melanogaster* was likely triggered by horizontal transfer from a different species. To identify the source of the horizontal transfer we investigated the long-read assemblies of 101 *Drosophila* species [Kim et al., 2021] and 99 insect species [Kim et al., 2021, Hotaling et al., 2021] (Supplementary table S7). Apart from *D. melanogaster* we found insertions with a high similarity to *Spoink* in *D. sechellia*, and species of the *willistoni* group, in particular *D. willistoni* (fig. 5A). *Spoink* insertions with a somewhat smaller similarity were found in *D. cardini* and *D. repleta*. No sequences similar to *Spoink* were found in the 99 insect species (supplementary fig. S6). To further shed light on the origin of the *Spoink* invasion we constructed a phylogenetic tree with full-length insertions of *Spoink* in *D. melanogaster, D. sechellia, D. cardini* and species of the *willistoni* group (fig. 5B). We did not find a full-length insertion of *Spoink* in *D. repleta*. This tree reveals that *Spoink* insertions in *D. sechellia* have very short branches, thus we suggest that the invasion in *D. sechellia* is also of recent origin.

**Figure 5.**
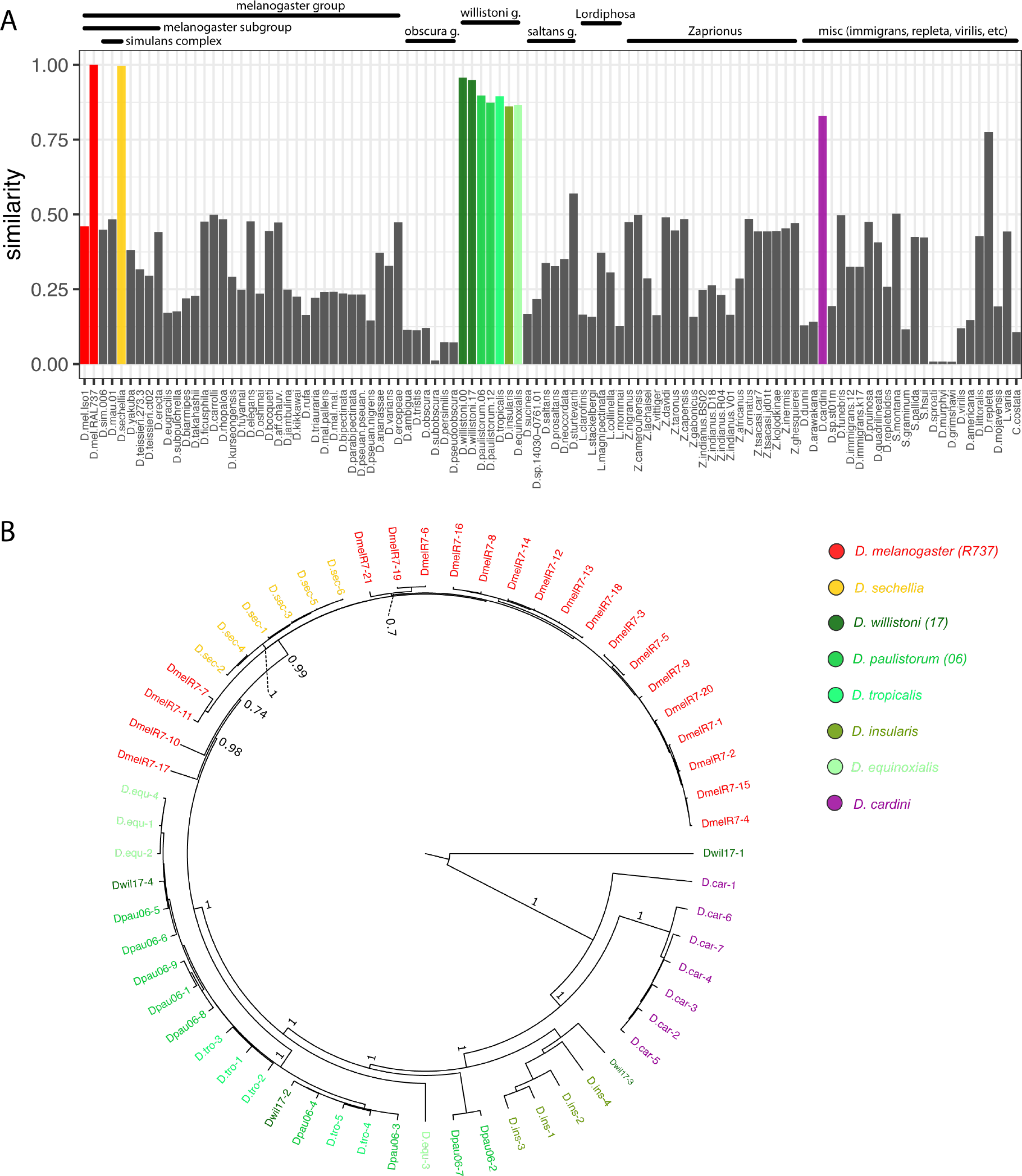
The *Spoink* invasion in *D. melanogaster* was likely triggered by a horizontal transfer from a species of the *willistoni* group. A) Similarity of TE insertions in long-read assemblies of diverse drosophilid species to *Spoink*. The barplots show for each species the similarity between *Spoink* and the best match in the assembly. For example, a value of 0.9 indicates that at least one insertion in an assembly has a high similarity (≈ 90%) to the consensus sequence of *Spoink*. B) Bayesian tree of *Spoink* insertions in the different drosophilid species. Only full-length insertions of *Spoink* (*>* 80% of the length) were considered. Note that *Spoink* insertions of *D. sechellia* are nested in insertions of *D. melanogaster*, while *Spoink* insertions of *D. melanogaster* are nested in insertions from the *willistoni* group (green shades).

However, *Spoink* insertions in *D. melanogaster* are nested within insertions from species of the *willistoni* group (fig. 5B). Our data thus suggest that, similar to the *P*-element invasion in *D. melanogaster* [Daniels et al., 1990], the *Spoink* invasion in *D. melanogaster* was also triggered by horizontal transfer from a species of the *willistoni* group. Species of the *willistoni* group are Neotropical, occurring throughout Central and South America [Burla et al., 1949, Spassky et al., 1971, Zanini et al., 2015]. Therefore horizontal transfer of *Spoink* only became feasible after *D. melanogaster* extended its habitat into the Americas approximately 200 years ago [A.H.Sturtevant, 1921, Johnson, 1913, Bock and Parsons, 1981]. Insertions of *D. cardini* are also nested within species of the *willistoni* group, suggesting that *D. cardini* also acquired *Spoink* by horizontal transfer from the *willistoni* group, likely independent of *D. melanogaster* (fig. 5B). *D. cardini* is also a Neotropical species and its range overlaps many species of the *willistoni* group, thus horizontal transfer between the species is physically feasible [Heed and Russell, 1971, Brisson et al., 2006].

In summary, similarly to the *P*-element, horizontal transfer from a species of the *willistoni* group likely triggered the *Spoink* invasion in *D. melanogaster*.

## Discussion

Here we showed that the LTR-retrotransposon *Spoink* invaded *D. melanogaster* populations between 1983 and 1993, after the spread of the *P*-element. Similarly to the *P*-element, the *Spoink* invasion was likely triggered by horizontal transfer from a species in the *willistoni* group.

The abundance of sequencing data from strains collected at different time points during the last century allowed us to pinpoint the timing of the invasion in a way that would not have been previously possible. *Spoink* appears to have rapidly spread throughout global populations of *D. melanogaster* between 1983 and 1993. The narrow time-window of 10 years is plausible as studies monitoring *P*-element invasions in experimental populations showed that the *P*-element can invade populations within 20-60 generations [Kofler et al., 2018, 2022, Selvaraju et al., 2022]. Assuming that natural *D. melanogaster* populations have about 15 generation per year [Pool, 2015], a TE could penetrate a natural *D. melanogaster* population within 1 - 3 years. Given this potential rapidness of TE invasions it is likely that *Spoink* spread quickly between 1983 and 1993. Since there is a gap between strains sampled at 1983 and 1993 we cannot further narrow down the timing of the invasion. Furthermore, the strains used for timing the invasions were sampled from diverse geographic regions and *Spoink* likely spread at different times in different geographic regions. If horizontal transfer from a *willistoni* species triggered the invasion, as suggested by our data, than *Spoink* will have first spread in *D. melanogaster* populations from South America (the habitat of *willistoni* species), followed by populations from North America and the other continents. Unfortunately we cannot infer the timing of the geographic spread of the *Spoink* invasion in different continents as *D. melanogaster* strains were not sampled sufficiently densely from different regions. Our work thus highlights the importance of efforts such as DrosEU, GDL and DrosRTEC to densely sample *Drosophila* strains in time and space [Kapun et al., 2020, Grenier et al., 2015, Machado et al., 2021]. It is also interesting to ask as to which extend human activity (e.g. trafficking of goods) contributed to the rapid spread of *Spoink*. Given that our analysis of the *Spoink* composition shows that geographically distant populations (Bejing/Ithaca or Netherlands/Zimbabwe) cluster together, human activity may have played a role. Increasing human activity could also explain why *Spoink* (invasion 1983-1993) seems to have spread faster than the *P*-element (1950-1980).

Our investigation of *Spoink* insertions in different drosophilid species suggests that the *Spoink* invasion in *D. melanogaster* was triggered by horizontal transfer from a species of the *willistoni* group. Although it is possible that we did not analyse the true donor species, we consider it unlikely to be a species outside of the *willistoni* group given the wide distribution of *Spoink* in all species in the *willistoni* group. In addition, the phylogenetic tree of *Spoink* has deep branches within the *willistoni* group, suggesting that *Spoink* is ancestral in this group. A related open question is when *Spoink* first entered *D. melanogaster* populations. Since a TE may initially solely spread in some isolated subpopulations there could be a considerable lag time between the horizontal transfer of a TE and its spread in worldwide population. Nevertheless, the horizontal transfer of *Spoink* must have happened between the spread of *D. melanogaster* into the habitat of the *willistoni* group, about 200 years ago, and the invasion of *Spoink* in worldwide populations between 1983 and 1993. In addition to the *P*-element, *Spoink* is the second TE that invaded *D. melanogaster* populations following horizontal transfer from a species of the *willistoni* group. Species from the *willistoni* group are very distantly related with *D. melanogaster* (about 100my [Obbard et al., 2012]) and we were thus wondering whether it is a coincidence that a species of the *willistoni* group is again acting as donor of a TE invasion in *D. melanogaster*. The recent habitat expansion of *D. melanogaster* into the Americas resulted in novel contacts with many species, in addition to species of the *willistoni* group, that might have acted as donors of novel TEs such as *D. pseudoobscura* or *D. persimilis*. Apart from mere chance, there are several feasible possibilities for this observation. First, it is feasible TEs of the *willistoni* group are exceptionally compatible with *D. melanogaster* at a molecular level. Second, some parasites targeting both *D. melanogaster* and species of the *willistoni* group could be efficient vectors for horizontal transfer of TEs. Third, the physical contact between *D. melanogaster* and some species of the *willistoni* group might be unusually tight, facilitating horizontal transfer of TEs by an unknown vector. Fourth, species of the *willistoni* group might be excepionally numerous resulting in elevated probability for horizontal transfer of a TE.

The *Spoink* invasion is the eighth identified TE invasion in *D. melanogaster* during the last two hundred years. As we argued previously, such a high rate of TE invasions is likely unusual during the evolution of the *D. melanogaster* lineage since the number of TE families in *D. melanogaster* is much smaller than what would be expected if this rate of invasions would persist [Scarpa et al., 2023]. One possible explanation for this high rate of TE invasions during the last two hundred years is that human activity contributed to the habitat expansion of *D. melanogaster*. Due to this habitat expansion *D. melanogaster* spread into the habitat of *D. willistoni* which enabled the horizontal transfer of *Spoink*. This raises the possibility that other species with recent habitat expansions also experienced unusually high rates of TE invasions. It is also interesting to ask whether the rate of TE invasions differs among species. For example cosmopolitan species, such as *D. melanogaster*, may generally experience higher rates of horizontal transfer than more locally confined species. The cosmopolitan distribution will bring species into contact with many diverse species, thereby increasing the opportunities for horizontal transfer of a TE.

The *Spoink* invasions also opens up several novel opportunities for research. First, the broad availability of strains with and without *Spoink* will enable testing whether *Spoink* activity induces phenotypic effects, similarly to hybrid dysgenesis described for the *P*-element, *I*-element and *hobo*, but not for *Tirant* [Bucheton et al., 1976, Kidwell et al., 1977, Blackman et al., 1987, Schwarz et al., 2021]. Second, it will be interesting to investigate whether some *Spoink* insertions participated in rapid adaptation of *D. melanogaster* populations, similar to a *P*-element insertion which contribute to insecticide resistance [Schmidt et al., 2010]. Third, it will enable studying *Spoink* invasions in experimental populations, shedding light on the dynamics of TE invasions, much as other recent studies investigating the invasion dynamics of the *P*-element [Kofler et al., 2018, 2022, Wang et al., 2023]. Fourth, investigation into the distribution of species that have been infected with *Spoink* will shed light on the networks of horizontal transfer in drosophild species. Fifth, the *Spoink* invasion provides an opportunity to study the establishment of the piRNA based host defence [similar to [Zhang et al., 2020, Selvaraju et al., 2022]]. For example we found that none of the piRNA cluster insertions are shared between individuals, suggesting there is no or solely weak selection for piRNA cluster insertions. Furthermore we found an extremely heterogeneous abundance of *Spoink* insertions in piRNA clusters where we could not find a single cluster insertions of *Spoink* in several strains. It is an important open question whether such a heterogeneous distribution is compatible with the trap model [Wierzbicki and Kofler, 2023]. One possibility is that a few cluster insertions in populations are sufficient to trigger the paramutation of regular *Spoink* insertions into piRNA producing loci [Scarpa and Kofler, 2023, Le Thomas et al., 2014b, Hermant et al., 2015]. These paramutated *Spoink* insertions may than compensate for the low number of *Spoink* insertions in piRNA-clusters [Scarpa and Kofler, 2023]. Paramutations could thus explain why several studies found that stand-alone insertions of TEs can nucleate their own piRNA production [Shpiz et al., 2014, Mohn et al., 2014, Wierzbicki and Kofler, 2023, Srivastav et al., 2023].

The war between transposons and their hosts is constantly raging, with potentially large fitness effects for the individuals in populations. Over the last two hundred years there have been at least eight invasions of TEs into *D. melanogaster*, each of which could disrupt fertility for example by inducing some form of hybrid dysgenesis. TEs are responsible for *>* 80% of visible spontaneous mutations in *D. melanogaster*, and produce more variation than all SNPs combined [Ashburner and Bergman, 2005, Green, 1988, Sankaranarayanan, 1988]. In the long read assemblies considered here, more than half of insertions of *Spoink* were into genes [Rech et al., 2022]. The recent *Spoink* invasion could thus have a significant impact on the evolution of *D. melanogaster* lineage.

## Materials and Methods

### Discovery of the recent *Spoink* invasion

We identified TE insertions in different long-read assemblies using RepeatMasker [Smit et al., 1996-2010] and the TE library from Chakraborty et al. [2021]. When comparing the TE composition between strains collected in the 1950’s and 1960’s [King et al., 2012, Chakraborty et al., 2019] and more recently collected strains (≥ 2003 [Rech et al., 2022] we noticed an element labeled ‘gypsy-7 DEl’ which was only present in short degraded copies in the older genomes but was present in full length copies in the more recent genomes (Supplementary table S1).

### Characterisation of *Spoink*

To generate a consensus sequence of *Spoink* we extracted the sequence of full-length matches of ‘*gypsy-7 DEl*’ plus some flanking sequences from long-read assemblies [Ten-15, RAL91, RAL176, RAL732, RAL737, Sto-22; [Rech et al., 2022]] and made a consensus sequence by performing multiple sequence alignment (MSA) with MUSCLE (v3.8.1551) [Edgar, 2004] and then choosing the most abundant nucleotide in each position of the MSA with a custom Python script (MSA2consensus).

The consensus sequence of the LTR was used to identify the TSD with our new tool LTRtoTE (https://github.com/Almo96/LTRtoTE). We used LTRdigest to identify the PBS of Spoink [Steinbiss et al., 2009]. We picked several sequences from each of the known LTR superfamily/groups using the consensus sequences of known TEs [Kapitonov and Jurka, 2003, Quesneville et al., 2005]. We performed a blastx search against the NCBI database to identify the RT domain in the consensus sequences of the TE [Wheeler et al., 2007]. We then performed a multiple sequence alignment of the amino-acid sequences of the RT domain using MUSCLE (v3.8.1551)[Edgar, 2004]. We obtained the xml file using BEAUti2[Bouckaert et al., 2019] (v2.7.5) and generated the trees with BEAST (v2.7.5)[Bouckaert et al., 2019]. The maximum credibility tree was built using TreeAnnotator (v2.7.5)[Bouckaert et al., 2019] and visualized with FigTree (v1.4.4, http://tree.bio.ed.ac.uk/software/figtree/).

### Distribution of *Spoink* insertions

Genes were annotated in each of the 31 genomes from [Rech et al., 2022] using the annotation of the reference genome of *D. melanogaster* (6.49; Flybase) and liftoff 1.6.3 [Shumate and Salzberg, 2021, Gramates et al., 2022]. The 1kb regions upstream of each gene were classified as putative promotors. The location of canonical *D. melanogaster* piRNA clusters was determined using CUSCO, which lifts over the flanks of known clusters in a reference genome to locate the homologous region in a novel genome [Wierzbicki et al., 2021]. The location of *Spoink* insertions within genes or clusters was determined with bedtools intersect [Quinlan and Hall, 2010]. To determine if genic insertions were shared or independent, the sequence of the insertion was extracted from each genome along with an extra 1 kb of flanking sequence on each end. Insertions purportedly in the same gene were then aligned, and if the flanks aligned they were considered shared insertions. To determine if cluster insertions were shared the flanking TE regions were aligned using Manna, which aligns TE annotations rather than sequences, to determine if there was any shared synteny in the surrounding TEs [Wierzbicki et al., 2023].

### Abundance of *Spoink* insertions in different *D. melanogaster* strains

We investigated the abundance of *Spoink* in multiple publicly available short-read data sets [Grenier et al., 2015, Schwarz et al., 2021, Long et al., 2013, Lange et al., 2021, Rech et al., 2022]. These data include genomic DNA from 183 *D. melanogaster* strains sampled at different geographic locations during the last centuries. For an overview of all analysed short-read data see Supplementary table S4. We mapped the short reads to a database consisting of the consensus sequences of TEs [Quesneville et al., 2005], the sequence of *Spoink* and three single copy genes (*rhino, trafficjam, rpl32*) with bwa bwasw (version 0.7.17-r1188) [Li and Durbin, 2009]. We used DeviaTE (v0.3.8)[Weilguny and Kofler, 2019] to estimate the abundance of *Spoink*. DeviaTE estimates the copy number of a TE (e.g. *Spoink*) by normalizing the coverage of the TE by the coverage of the single copy genes. We also used DeviaTE to visualize the abundance and diversity of *Spoink* as well as to compute the frequency of SNPs in *Spoink* (see below).

To identify *Spoink* insertions in 49 long-read assemblies of *D. melanogaster* strains collected during the last 100 years we used RepeatMasker [Smit et al., 1996-2010] (open-4.0.7; -no-is -s -nolow). For an overview of all analysed assemblies see Supplementary table S5[Chakraborty et al., 2019, Wierzbicki et al., 2021, Hoskins et al., 2015, Rech et al., 2022]. For estimating the abundance of *Spoink* in the long-read assemblies we solely considered canonical *Spoink* insertions (*>* 80% of length, *<* 5% sequence divergence).

### Population frequency of *Spoink* insertions

For every putative *Spoink* insertion (including degraded ones) in the eight long-read assemblies of individuals from Raleigh [Rech et al., 2022], we extracted the sequence of the insertion plus 1 kb of flanking sequence with bedtools [Quinlan and Hall, 2010]. The sequence of the *Spoink* insertion was removed with seqkit [Shen et al., 2016] and the flanking sequences were mapped to the *AKA017* genome (i.e. the common coordinate system) with minimap2 allowing for spliced mappings [Li, 2018, Shen et al., 2016, Rech et al., 2022]. The mapping location of each read was extracted and if they overlapped between strains they were considered putative shared sites. Regions with overlapping reads were visually inspected in IGV (v2.4.14) and if the mapping location was shared they were considered shared insertions sites [Robinson et al., 2010, Thorvaldsdóttir et al., 2012].

### PCR

To validate whether *Spoink* is absent in old *D. melanogaster* strains but present in recent strains we used PCR. We designed two primers pairs for *Spoink* and one for *vasa* as a control. We extracted DNA from different strains of *D. melanogaster* (*Lausanne-S, Hikone-R,iso-1,RAL59, RAL176, RAL737*) using a high salt extraction protocol [Miller et al., 1988]. We designed two primers pairs for *Spoink* (P1,P2) and one for the gene *Vasa* (P1 FWD TCAGAAGTGGGATCGGGCTCGG, P1 REV CAGTAGAGCACCATGCCGACGC, P2 FWD ATGGACCGTAATGGCAGCAGCG, P2 REV ACACTCCGCGCCAGAGTCAAAC, Vasa FWD AACGAGGCGAGGAAGTTTGC, Vasa REV GCGATCACTACATGGCAGCC) We used the following PCR conditions: 1 cycle of 95°C for 3 minutes; 33 cycles of 95°C for 30 seconds, 58°C for 30 seconds and 72°C for 20 seconds; 1 cycle of 72°C for 6 minutes.

### Small RNAs

To identify piRNAs complementary to *Spoink* we analysed the small-RNA data from 10 GDL strains [Luo et al., 2020]. The adaptor sequence GAATTCTCGGGTGCCAAGG was removed using cutadapt (v4.4 [Martin, 2011]). We filtered for reads having a length between 18 and 36nt and aligned the reads to a database consisting of *D. melanogaster* miRNAs, mRNAs, rRNAs, snRNAs, snoRNAs, tRNAs [Thurmond et al., 2019], and TE sequences [Quesneville et al., 2005] with novoalign (v3.09.04). We used previously developed Python scripts [Selvaraju et al., 2022] to compute ping-pong signatures and to visualize the piRNA abundance along the sequence of *Spoink*.

### UMAP

We used the frequencies of SNPs in the sequence of *Spoink* to compute the UMAP. This frequencies reflect the *Spoink* composition in a given sample. For example if a specimen has 20 *Spoink* insertions and a biallelic SNP with a frequency of 0.8 at a given site in *Spoink* than about 16 *Spoink* insertions will have the SNP and 4 will not have it. The frequency of the *Spoink* SNPs was estimated with DeviaTE [Weilguny and Kofler, 2019]. Solely bi-allelic SNPs were used and SNPs only found in few samples were removed (≤ 3 samples) UMAPs were created in R (umap package; v0.2.10.0 [McInnes et al., 2018]).

### Origin of horizontal transfer

To identify the origin of the horizontal transfer of *Spoink* we used RepeatMasker [Smit et al., 1996-2010] (open-4.0.7; -no-is -s -nolow) to identify sequences with similarity to *Spoink* in the long-read assemblies of 101 drosophilid species and in 99 different insect species [Kim et al., 2021, Hotaling et al., 2021] (Supplementary table S7). We included the long-read assembly of the *D. melanogaster* strain *RAL737* in the analysis [Rech et al., 2022]. We used a Python script to identify in each assembly the best hit with *Spoink* (i.e. the highest alignment score) and than estimated the similarity between this best hit and *Spoink*. The similarity was computed as *s* = *rms*_*best*_*/rms*_*max*_, where *rms*_*best*_ is the highest RepeatMasker score (rms) in a given assembly and *rms*_*max*_ the highest score in any of the analysed assemblies. A *s* = 0 indicates no similarity to the consensus sequence of *Spoink* whereas *s* = 1 represent the highest possible similarity. To generate a phylogenetic tree we identified *Spoink* insertions in the assemblies of the 101 drosophilid species and *RAL737* using RepeatMasker. We extracted the sequences of full-length insertions (*>* 80% of the length) from species having at least one full-length insertion using bedtools [Quinlan and Hall, 2010](v2.30.0). A multiple sequence alignment of the *Spoink* insertions was generated with MUSCLE (v3.8.1551)[Edgar, 2004] and a tree was generated with BEAST (v2.7.5)[Bouckaert et al., 2019].

## Acknowledgments

We thank Matthew Beaumont for the idea to call the here described transposon *Spoink*. We thank Neda Barghi and Claudia Ramirez Lanzas for providing fly strains used for PCR. SS would like to thank J. B. Signor for helpful comments on the manuscript. RK, RP, and AS thank all members of the Institute of Population Genetics for feedback and support.

## Author contributions

SS discovered *Spoink*. RK and SS conceived the study. RP, AS, SS and RK analysed the data and generated the figures. AS performed PCR. RK and SS wrote the first draft. RP and AS contributed to writing. PN assisted with data collection.

## Data Availability

The consensus sequence of *Spoink* as well as the sequences of the six PCR amplicons are available at https://github.com/rpianezza/Dmel-Spoink/tree/main/releasedseqs. The tool LTRtoTE is available on GitHub (https://github.com/Almo96/LTRtoTE). The analysis performed in this work have been documented with RMarkdown and have been made publicly available, together with the resulting figures, at GitHub (https://github.com/rpianezza/Dmel-Spoink; see *.md files).

## References

A.H. Sturtevant. The North American species of Drosophila. Nature, 107:1476–4687, 1921.

V. Andreev, C. Yu, and e. a. Wang, J. Panoramix SUMOylation on chromatin connects the piRNA pathway to the cellular heterochromatin machinery. Nat Struct Mol Biol, 29:130–142, 2022.

D. Anxolabéhère, D. Nouaud, G. Périquet, and P. Tchen. P-element distribution in Eurasian populations of Drosophila melanogaster : a genetic and molecular analysis. Proceedings of the National Academy of Sciences, 82(16):5418–5422, 1985.

D. Anxolabéhère, M. G. Kidwell, and G. Periquet. Molecular characteristics of diverse populations are consistent with the hypothesis of a recent invasion of Drosophila melanogaster by mobile P elements. Molecular Biology and Evolution, 5(3):252–269, 1988.

M. Ashburner and C. M. Bergman. Drosophila melanogaster : A case study of a model genomic sequence and its consequences. Cold Spring Harbor perspectives in biology, 15:1661–1667, 2005. doi: 10.1101/gr.3726705.15.

C. M. Bergman, H. Quesneville, D. Anxolabéhère, and M. Ashburner. Recurrent insertion and duplication generate networks of transposable element sequences in the Drosophila melanogaster genome. Genome Biology, 7(11):R112, 2006.

R. K. Blackman, R. Grimaila, M. Macy, D. Koehler, and W. M. Gelbart. Mobilization of hobo elements residing within the decapentaplegic gene complex: Suggestion of a new hybrid dysgenesis system in Drosophila melanogaster. Cell, 49(4):497–505, 1987.

J. P. Blumenstiel. Birth, school, work, death, and resurrection: The life stages and dynamics of transposable element proliferation. Genes, 10(5):336, 2019.

I. Bock and P. Parsons. Species of Australia and New Zealand. In M. Ashburner, L. Carson, and J. J. Thompson, editors, The genetics and biology of Drosophila, volume 3a, pages 349–393. Academic Press, Oxford, 1981.

E. Bonnivard and D. Higuet. Stability of European natural populations of Drosophila melanogaster with regard to the P-M system: a buffer zone made up of Q populations. Journal of Evolutionary Biology, 12 (4):633–647, 1999.

R. Bouckaert, T. G. Vaughan, J. Barido-Sottani, S. Duchêne, M. Fourment, A. Gavryushkina, J. Heled, G. Jones, D. Kühnert, N. D. Maio, M. Matschiner, F. K. Mendes, N. F. Müller, H. A. Ogilvie, L. du Plessis, A. Popinga, A. Rambaut, D. Rasmussen, I. Siveroni, M. A. Suchard, C.-H. Wu, D. Xie, C. Zhang, T. Stadler, and A. J. Drummond. BEAST 2.5: An advanced software platform for bayesian evolutionary analysis. PLOS Computational Biology, 15(4):e1006650, Apr. 2019.

J. Brennecke, A. A. Aravin, A. Stark, M. Dus, M. Kellis, R. Sachidanandam, and G. J. Hannon. Discrete small RNA-generating loci as master regulators of transposon activity in Drosophila. Cell, 128(6):1089–1103, 2007.

J. Brennecke, C. D. Malone, A. A. Aravin, R. Sachidanandam, A. Stark, and G. J. Hannon. An epigenetic role for maternally inherited piRNAs in transposon silencing. Science, 322(5906):1387–1392, 2008.

J. A. Brisson, J. Wilder, and H. Hollocher. Phylogenetic analysis of the cardini group of Drosophila with respect to changes in pigmentation. Evolution, 60(6):1228–1241, 2006.

A. Bucheton, J. Lavige, G. Picard, and P. L’heritier. Non-mendelian female sterility in Drosophila melanogaster : quantitative variations in the efficiency of inducer and reactive strains. Heredity, 36(3): 305–314, 1976.

H. Burla, A. B. da Cunha, A. R. Cordeiro, T. Dobzhansky, C. Malogolowkin, and C. Pavan. The Willistoni Group of Sibling Species of Drosophila. Evolution, 3(4):300–314, 1949.

M. Chakraborty, J. J. Emerson, S. J. Macdonald, and A. D. Long. Structural variants exhibit widespread allelic heterogeneity and shape variation in complex traits. Nature Communications, 10(1):4872, 2019.

M. Chakraborty, C. Chang, D. Khost, J. A. J Vedanayagam, Y. Liao, K. Montooth, C. Meiklejohn, A. Larracuente, and J. Emerson. Evolution of genome structure in the Drosophila simulans species complex. Genome Research, 31:380–396, 2021.

S. B. Daniels, K. R. Peterson, L. D. Strausbaugh, M. G. Kidwell, and A. Chovnick. Evidence for horizontal transmission of the P transposable element between Drosophila species. Genetics, 124(2):339–55, 1990.

A. Diaz-Papkovich, L. Anderson-Trocmé, and S. Gravel. A review of UMAP in population genetics. Journal of Human Genetics, 66(1):85–91, 2021.

R. C. Edgar. Muscle: multiple sequence alignment with high accuracy and high throughput. Nucleic acids research, 32(5):1792–1797, 2004.

D. G. Eickbush and T. H. Eickbush. Vertical transmission of the retrotransposable elements R1 and R2 during the evolution of the Drosophila melanogaster species subgroup. Genetics, 139(2):671–684, 1995.

T. H. Eickbush and H. S. Malik. Origins and evolution of retrotransposons, volume 93. ASM Press, 2002.

S. F. Elena, L. Ekunwe, N. Hajela, S. A. Oden, and R. E. Lenski. Distribution of fitness effects caused by random insertion mutations in Escherichia coli. Genetica, 102-103:349–358, 1998.

C. E. Ellison and W. Cao. Nanopore sequencing and Hi-C scaffolding provide insight into the evolutionary dynamics of transposable elements and piRNA production in wild strains of Drosophila melanogaster. Nucleic Acids Research, 48(1):1–14, 2020.

D. J. Finnegan. Eukaryotic transposable elements and genome evolution. Trends in Genetics, 5(4):103–107, 1989.

C. Goriaux, E. Théron, E. Brasset, and C. Vaury. History of the discovery of a master locus producing piRNAs: The flamenco/COM locus in Drosophila melanogaster. Frontiers in Genetics, 5:257, 2014.

L. S. Gramates, J. Agapite, H. Attrill, B. R. Calvi, M. A. Crosby, G. Dos Santos, J. L. Goodman, D. Goutte-Gattat, V. K. Jenkins, T. Kaufman, et al. Flybase: a guided tour of highlighted features. Genetics, 220 (4):iyac035, 2022.

M. M. Green. Mobile DNA elements and spontaneous gene mutation. Banbury Rep, 30:41–50, 1988.

J. K. Grenier, J. R. Arguello, M. C. Moreira, S. Gottipati, J. Mohammed, S. R. Hackett, R. Boughton, A. J. Greenberg, and A. G. Clark. Global diversity lines–a five-continent reference panel of sequenced Drosophila melanogaster strains. G3: Genes, Genomes, Genetics, 5(4):593–603, 2015.

L. S. Gunawardane, K. Saito, K. M. Nishida, K. Miyoshi, Y. Kawamura, T. Nagami, H. Siomi, and M. C. Siomi. A slicer-mediated mechanism for repeat-associated siRNA 5’ end formation in Drosophila. Science, 315(5818):1587–1590, 2007.

W. B. Heed and J. S. Russell. Phylogeny and population structure in island and continental species of the cardini group of Drosophila studied by inversion analysis. Stud Genet, 1971.

C. Hermant, A. Boivin, L. Teysset, V. Delmarre, A. Asif-Laidin, M. Van Den Beek, C. Antoniewski, and S. Ronsseray. Paramutation in Drosophila requires both nuclear and cytoplasmic actors of the piRNA pathway and induces cis-spreading of piRNA production. Genetics, 201(4):1381–1396, 2015.

R. A. Hoskins, J. W. Carlson, K. H. Wan, S. Park, I. Mendez, S. E. Galle, B. W. Booth, B. D. Pfeiffer, R. A. George, R. Svirskas, et al. The Release 6 reference sequence of the Drosophila melanogaster genome. Genome Research, 25(3):445–458, 2015.

S. Hotaling, J. S. Sproul, J. Heckenhauer, A. Powell, A. M. Larracuente, S. U. Pauls, J. L. Kelley, and P. B. Frandsen. Long Reads Are Revolutionizing 20 Years of Insect Genome Sequencing. Genome Biology and Evolution, 13(8):evab138, 06 2021.

C. Johnson. The distribution of some species of Drosophila. Psyche, 20:202–205, 1913.

V. V. Kapitonov and J. Jurka. Molecular paleontology of transposable elements in the Drosophila melanogaster genome. Proceedings of the National Academy of Sciences of the United States of America, 100(11):6569–74, 2003. ISSN 0027-8424.

M. Kapun, M. G. Barrón, F. Staubach, D. J. Obbard, R. A. W. Wiberg, J. Vieira, C. Goubert, O. Rota-Stabelli, M. Kankare, M. Bogaerts-Márquez, et al. Genomic analysis of European Drosophila melanogaster populations reveals longitudinal structure, continent-wide selection, and previously unknown DNA viruses. Molecular Biology and Evolution, 37(9):2661–2678, 2020.

M. G. Kidwell. Evolution of hybrid dysgenesis determinants in Drosophila melanogaster. Proceedings of the National Academy of Sciences, 80(6):1655–1659, 1983.

M. G. Kidwell, J. F. Kidwell, and J. A. Sved. Hybrid dysgenesis in Drosophila melanogaster : A syndrome of aberrant traits including mutations, sterility and male recombination. Genetics, 86(4):813–833, 1977.

B. Y. Kim, J. R. Wang, D. E. Miller, O. Barmina, E. Delaney, A. Thompson, A. A. Comeault, D. Peede, E. R. D’Agostino, J. Pelaez, J. M. Aguilar, D. Haji, T. Matsunaga, E. E. Armstrong, M. Zych, Y. Ogawa, M. Stamenković -Radak, M. Jelić, M. S. Veselinović, M. Tanasković, P. Erić, J.-J. Gao, T. K. Katoh, M. J. Toda, H. Watabe, M. Watada, J. S. Davis, L. C. Moyle, G. Manoli, E. Bertolini, V. Koštál, R. S. Hawley, A. Takahashi, C. D. Jones, D. K. Price, N. Whiteman, A. Kopp, D. R. Matute, and D. A. Petrov. Highly contiguous assemblies of 101 drosophilid genomes. eLife, 10:e66405, 2021.

E. G. King, C. M. Merkes, C. L. McNeil, S. R. Hoofer, S. Sen, K. W. Broman, A. D. Long, and S. J. Macdonald. Genetic dissection of a model complex trait using the Drosophila Synthetic Population Resource. Genome Research, 22(8):1558–1566, 2012.

R. Kofler. Dynamics of Transposable Element Invasions with piRNA Clusters. Molecular Biology and Evolution, 36(7):1457–1472, 2019.

R. Kofler, T. Hill, V. Nolte, A. Betancourt, and C. Schlötterer. The recent invasion of natural Drosophila simulans populations by the P-element. PNAS, 112(21):6659–6663, 2015.

R. Kofler, K.-A. Senti, V. Nolte, R. Tobler, and C. Schlötterer. Molecular dissection of a natural transposable element invasion. Genome Research, 28(6):824–835, 2018.

R. Kofler, V. Nolte, and C. Schlötterer. The Transposition Rate Has Little Influence on the Plateauing Level of the P-element. Molecular Biology and Evolution, 39(7):msac141, 2022.

J. D. Lange, H. Bastide, J. B. Lack, and J. E. Pool. A Population Genomic Assessment of Three Decades of Evolution in a Natural Drosophila Population. Molecular Biology and Evolution, 39(2), 2021.

A. Le Thomas, A. K. Rogers, A. Webster, G. K. Marinov, S. E. Liao, E. M. Perkins, J. K. Hur, A. A. Aravin, and K. F. Tóth. Piwi induces piRNA-guided transcriptional silencing and establishment of a repressive chromatin state. Genes and Development, 27(4):390–399, 2013.

A. Le Thomas, G. K. Marinov, and A. A. Aravin. A transgenerational process defines piRNA biogenesis in Drosophila virilis. Cell Reports, 8(6):1617–1623, 2014a.

A. Le Thomas, E. Stuwe, S. Li, J. Du, G. Marinov, N. Rozhkov, Y. C. A. Chen, Y. Luo, R. Sachidanandam, K. F. Toth, D. Patel, and A. A. Aravin. Transgenerationally inherited piRNAs trigger piRNA biogenesis by changing the chromatin of piRNA clusters and inducing precursor processing. Genes and Development, 28(15):1667–1680, 2014b.

H. Li. Minimap2: pairwise alignment for nucleotide sequences. Bioinformatics, 34(18):3094–3100, 2018.

H. Li and R. Durbin. Fast and accurate short read alignment with Burrows–Wheeler transform. Bioinformatics, 25(14):1754–1760, 2009.

D. H. Lindsley and E. H. Grell. Genetic variations of Drosophila melanogaster. Carnegie Institute of Washington Publication, 1968. ISBN 0099-4936.

R. S. Linheiro and C. M. Bergman. Whole genome resequencing reveals natural target site preferences of transposable elements in Drosophila melanogaster. PloS one, 7(2):e30008, 2012.

Q. Long, F. A. Rabanal, D. Meng, C. D. Huber, A. Farlow, A. Platzer, Q. Zhang, B. J. Vilhjálmsson, A. Korte, V. Nizhynska, et al. Massive genomic variation and strong selection in Arabidopsis thaliana lines from Sweden. Nature genetics, 45(8):884–890, 2013.

E. L. S. Loreto, C. M. A. Carareto, and P. Capy. Revisiting horizontal transfer of transposable elements in Drosophila. Heredity, 100(6):545–54, 2008.

S. Luo, H. Zhang, Y. Duan, X. Yao, A. G. Clark, and J. Lu. The evolutionary arms race between transposable elements and piRNAs in Drosophila melanogaster. BMC Evolutionary Biology, 20(1):1–18, 2020.

H. E. Machado, A. O. Bergland, R. Taylor, S. Tilk, E. Behrman, K. Dyer, D. K. Fabian, T. Flatt, J. González, T. L. Karasov, et al. Broad geographic sampling reveals the shared basis and environmental correlates of seasonal adaptation in drosophila. Elife, 10:e67577, 2021.

C. D. Malone and G. J. Hannon. Small RNAs as Guardians of the Genome. Cell, 136(4):656–668, 2009.

M. Martin. Cutadapt removes adapter sequences from high-throughput sequencing reads. EMBnet. journal, 17(1):10–12, 2011.

L. McInnes, J. Healy, and J. Melville. Umap: Uniform manifold approximation and projection for dimension reduction. arXiv preprint arXiv:1802.03426, 2018.

S. A. Miller, D. D. Dykes, and H. F. Polesky. A simple salting out procedure for extracting DNA from human nucleated cells. Nucleic acids research, 16(3):1215, 1988.

F. Mohn, G. Sienski, D. Handler, and J. Brennecke. The rhino-deadlock-cutoff complex licenses noncanonical transcription of dual-strand piRNA clusters in Drosophila. Cell, 157(6):1364–1379, 2014.

J. Nelson, A. Slicko, and Y. Yamashita. The retrotransposon R2 maintains Drosophila ribosomal DNA repeats. PNAS, 120(23):e2221613120, 2023.

D. J. Obbard, J. Maclennan, K.-W. Kim, A. Rambaut, P. M. O’Grady, and F. M. Jiggins. Estimating divergence dates and substitution rates in the Drosophila phylogeny. Molecular Biology and Evolution, 29 (11):3459–3473, 2012.

E. Pasyukova, S. Nuzhdin, T. Morozova, and T. Mackay. Accumulation of transposable elements in the genome of Drosophila melanogaster is associated with a decrease in fitness. Journal of Heredity, 95(4): 284–290, 2004.

J. Peccoud, V. Loiseau Cordaux, and C. Gilbert. Massive horizontal transfer of transposable elements in insects. Proc Natl Acad Sci U S A, 114(18):4721–26, 2017.

G. Periquet, M. H. Hamelin, Y. Bigot, and A. Lepissier. Geographical and historical patterns of distribution of hobo elements in Drosophila melanogaster populations. Journal of Evolutionary Biology, 2(3):223–229, 1989.

J. E. Pool. The Mosaic Ancestry of the Drosophila Genetic Reference Panel and the D. melanogaster Reference Genome Reveals a Network of Epistatic Fitness Interactions. Molecular biology and evolution, page msv194, 2015.

H. Quesneville, C. M. Bergman, O. Andrieu, D. Autard, D. Nouaud, M. Ashburner, and D. Anxolabéhère. Combined evidence annotation of transposable elements in genome sequences. PLoS Computational Biology, 1(2):166–175, 2005.

A. R. Quinlan and I. M. Hall. BEDTools: a flexible suite of utilities for comparing genomic features. Bioinformatics (Oxford, England), 26(6):841–842, 2010.

P. Rangan, C. D. Malone, C. Navarro, S. P. Newbold, P. S. Hayes, R. Sachidanandam, G. J. Hannon, and R. Lehmann. piRNA production requires heterochromatin formation in Drosophila. Current Biology, 21 (16):1373–1379, 2011.

G. E. Rech, S. Radío, S. Guirao-Rico, L. Aguilera, V. Horvath, L. Green, H. Lindstadt, V. Jamilloux, H. Quesneville, and J. González. Population-scale long-read sequencing uncovers transposable elements associated with gene expression variation and adaptive signatures in drosophila. Nature Communications, 13(1):1948, 2022.

M. D. Robinson, D. J. McCarthy, and G. K. Smyth. edgeR: a Bioconductor package for differential expression analysis of digital gene expression data. Bioinformatics, 26(1):139–140, 2010.

K. Sankaranarayanan. Mobile genetic elements, spontaneous mutations, and the assessment of genetic radiation hazards in man. Cold Spring Harbor, 1988.

P. Sarkies, M. E. Selkirk, J. T. Jones, V. Blok, T. Boothby, B. Goldstein, B. Hanelt, A. Ardila-Garcia, N. M. Fast, P. M. Schiffer, C. Kraus, M. J. Taylor, G. Koutsovoulos, M. L. Blaxter, and E. A. Miska. Ancient and novel small RNA pathways compensate for the loss of piRNAs in multiple independent nematode lineages. PLoS Biol., 13(2):1–20, 2015.

A. Scarpa and R. Kofler. The impact of paramutations on the invasion dynamics of transposable elements. Genetics, page iyad181, 2023.

A. Scarpa, R. Pianezza, F. Wierzbicki, and R. Kofler. Genomes of historical specimens reveal multiple invasions of ltr retrotransposons in drosophila melanogaster populations during the 19th century. bioRxiv, 2023. doi: 10.1101/2023.06.06.543830.

S. Schaack, C. Gilbert, and C. Feschotte. Promiscuous DNA: horizontal transfer of transposable elements and why it matters for eukaryotic evolution. Trends in ecology & evolution, 25(9):537–46, 2010.

J. M. Schmidt, R. T. Good, B. Appleton, J. Sherrard, G. C. Raymant, M. R. Bogwitz, J. Martin, P. J. Daborn, M. E. Goddard, P. Batterham, et al. Copy number variation and transposable elements feature in recent, ongoing adaptation at the cyp6g1 locus. PLoS genetics, 6(6):e1000998, 2010.

P. S. Schnable, S. Pasternak, C. Liang, J. Zhang, L. Fulton, T. A. Graves, P. Minx, A. D. Reily, L. Courtney, S. S. Kruchowski, C. Tomlinson, C. Strong, K. Delehaunty, C. Fronick, B. Courtney, S. M. Rock, E. Belter, F. Du, K. Kim, R. M. Abbott, M. Cotton, A. Levy, P. Marchetto, K. Ochoa, S. M. Jackson, B. Gillam, W. Chen, L. Yan, J. Higginbotham, M. Cardenas, J. Waligorski, E. Applebaum, L. Phelps, J. Falcone, K. Kanchi, T. Thane, A. Scimone, N. Thane, J. Henke, T. Wang, J. Ruppert, N. Shah, K. Rotter, J. Hodges, E. Ingenthron, M. Cordes, S. Kohlberg, J. Sgro, B. Delgado, K. Mead, A. Chinwalla, S. Leonard, K. Crouse, K. Collura, D. Kudrna, J. Currie, R. He, A. Angelova, S. Rajasekar, T. Mueller, R. Lomeli, G. Scara, A. Ko, K. Delaney, M. Wissotski, G. Lopez, D. Campos, M. Braidotti, E. Ashley, W. Golser, H. Kim, S. Lee, J. Lin, Z. Dujmic, W. Kim, J. Talag, A. Zuccolo, C. Fan, A. Sebastian, M. Kramer, L. Spiegel, L. Nascimento, T. Zutavern, B. Miller, C. Ambroise, S. Muller, W. Spooner, A. Narechania, L. Ren, S. Wei, and S. Kumari. The B73 Maize Genome: Complexity, Diversity, and Dynamics. Science, 326(5956):1112–1115, 2009.

F. Schwarz, F. Wierzbicki, K.-A. Senti, and R. Kofler. Tirant Stealthily Invaded Natural Drosophila melanogaster Populations during the Last Century. Molecular Biology and Evolution, 38(4):1482–1497, 2021.

D. Selvaraju, F. Wierzbicki, and R. Kofler. P-element invasions in Drosophila erecta shed light on the establishment of host control over a transposable element. bioRxiv, 2022.

W. Shen, S. Le, Y. Li, and F. Hu. Seqkit: A cross-platform and ultrafast toolkit for fasta/q file manipulation. PLOS ONE, 11(10):1–10, 10 2016.

S. Shpiz, S. Ryazansky, I. Olovnikov, Y. Abramov, and A. Kalmykova. Euchromatic transposon insertions trigger production of novel pi-and endo-siRNAs at the target sites in the Drosophila germline. PLoS Genetics, 10(2):e1004138, 2014.

A. Shumate and S. Salzberg. Liftoff: accurate mapping of gene annotations. Bioinformatics, 37(12):1639–1643, 2021.

G. Sienski, D. Dönertas, and J. Brennecke. Transcriptional silencing of transposons by Piwi and maelstrom and its impact on chromatin state and gene expression. Cell, 151(5):964–980, 2012.

S. Signor, J. Vedanayagam, B. Kim, F. Wierzbicki, R. Kofler, and E. Lai. Rapid evolutionary diversification of the flamenco locus across simulans clade drosophila species. PLoS Genet, 19, 2023.

G. Simmons. Horizontal transfer of hobo transposable elements within the Drosophila melanogaster species complex: evidence from dna sequencing. Molecular biology and evolution, 9(6):1050–1060, 1992.

F. A. Smit, R. Hubley, and P. Green. RepeatMasker Open-3.0, 1996-2010. URL http://www.repeatmasker.org.

Spassky, R. C. Richmond, S. Perez-Salas, O. Pavlovsky, C. A. Mourao, A. S. Hunter, H. Hoenigsberg, T. Dobzhansky, and F. J. Ayala. Geography of the Sibling Species Related to Drosophila willistoni, and of the Semispecies of the Drosophila Paulistorum Complex. Evolution, 25(1):129–143, 1971.

S. Srivastav, C. Feschotte, and A. G. Clark. Rapid evolution of piRNA clusters in the Drosophila melanogaster ovary. bioRxiv, 2023.

S. Steinbiss, U. Willhoeft, G. Gremme, and S. Kurtz. Fine-grained annotation and classification of de novo predicted LTR retrotransposons. Nucleic acids research, 37(21):7002–7013, 2009.

H. Thorvaldsdóttir, J. T. Robinson, and J. P. Mesirov. Integrative Genomics Viewer (IGV): high-performance genomics data visualization and exploration. Briefings in bioinformatics, 2012.

J. Thurmond, J. L. Goodman, V. B. Strelets, H. Attrill, L. S. Gramates, S. J. Marygold, B. B. Matthews, G. Millburn, G. Antonazzo, V. Trovisco, et al. Flybase 2.0: the next generation. Nucleic acids research, 47(D1):D759–D765, 2019.

L. Wang, S. Zhang, S. Hadjipanteli, L. Saiz, L. Nguyen, E. Silva, and E. Kelleher. P-element invasion fuels molecular adaptation in laboratory populations of Drosophila melanogaster. Evolution, 77(4):980–994, 2023.

L. Weilguny and R. Kofler. DeviaTE: Assembly-free analysis and visualization of mobile genetic element composition. Molecular Ecology Resources, 19(5):1346–1354, 2019.

D. L. Wheeler, T. Barrett, D. A. Benson, S. H. Bryant, K. Canese, V. Chetvernin, D. M. Church, M. DiCuccio, R. Edgar, S. Federhen, et al. Database resources of the national center for biotechnology information. Nucleic acids research, 35(suppl 1):D5–D12, 2007.

T. Wicker, F. Sabot, A. Hua-Van, J. L. Bennetzen, P. Capy, B. Chalhoub, A. Flavell, P. Leroy, M. Morgante, O. Panaud, et al. A unified classification system for eukaryotic transposable elements. Nature Reviews Genetics, 8(12):973–982, 2007.

F. Wierzbicki and R. Kofler. The composition of piRNA clusters in Drosophila melanogaster deviates from expectations under the trap model. BMC Biology, 2023.

F. Wierzbicki, F. Schwarz, O. Cannalonga, and R. Kofler. Novel quality metrics allow identifying and generating high-quality assemblies of piRNA clusters. Molecular Ecology Resources, 2021. doi: 10.1111/1755-0998.13455.

F. Wierzbicki, R. Kofler, and S. Signor. Evolutionary dynamics of piRNA clusters in Drosophila. Molecular Ecology, 32(6):1306–1322, 2023.

S. Yamanaka, M. C. Siomi, and H. Siomi. piRNA clusters and open chromatin structure. Mobile DNA, 5(1): 22, 2014.

R. Zanini, M. Deprá, and V. Valante. On the geographic distribution of the Drosophila willistoni group (Diptera, Drosophilidae) - updated geographic distribution of the Neotropical willistoni subgroup. 98: 39–43, 01 2015.

V. Zanni, A. Eymery, M. Coiffet, M. Zytnicki, I. Luyten, H. Quesneville, C. Vaury, and S. Jensen. Distribution, evolution, and diversity of retrotransposons at the flamenco locus reflect the regulatory properties of piRNA clusters. Proceedings of the National Academy of Sciences, 110(49):19842–19847, 2013.

S. Zhang, B. Pointer, and E. S. Kelleher. Rapid evolution of piRNA-mediated silencing of an invading transposable element was driven by abundant de novo mutations. Genome Research, 30(4):566–575, 2020.

