## Supplementary Material for "Spoink, a LTR retrotransposon, invaded *D. melanogaster* populations in the 1990s"

### Supplementary tables and figures

October 30, 2023

#### Supplementary figures

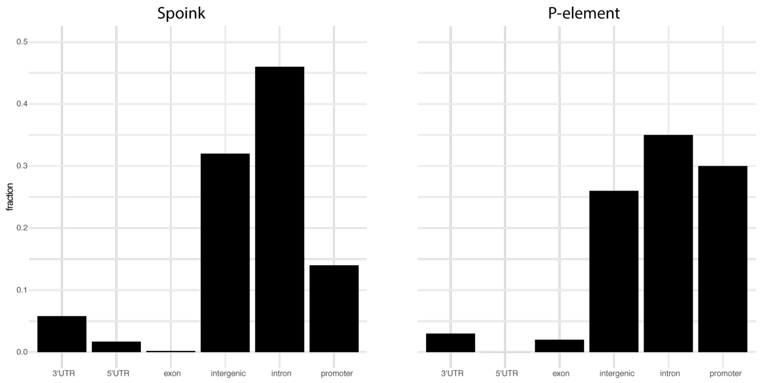


Figure 1: Abundance of *Spoink* and P-element insertions in different genomic features. TE insertions were identified in 31 long-read assemblies of *D. melanogaster* [Rech et al., 2022] and the reference annotation was lifted to each assembly with liftoff [Shumate and Salzberg, 2021, Gramates et al., 2022]. Note that the P-element has a pronounced insertion bias in promoters (defined as 1000bp upstream of the first exon) whereas *Spoink* insertions are largely found in introns and intergenic regions.

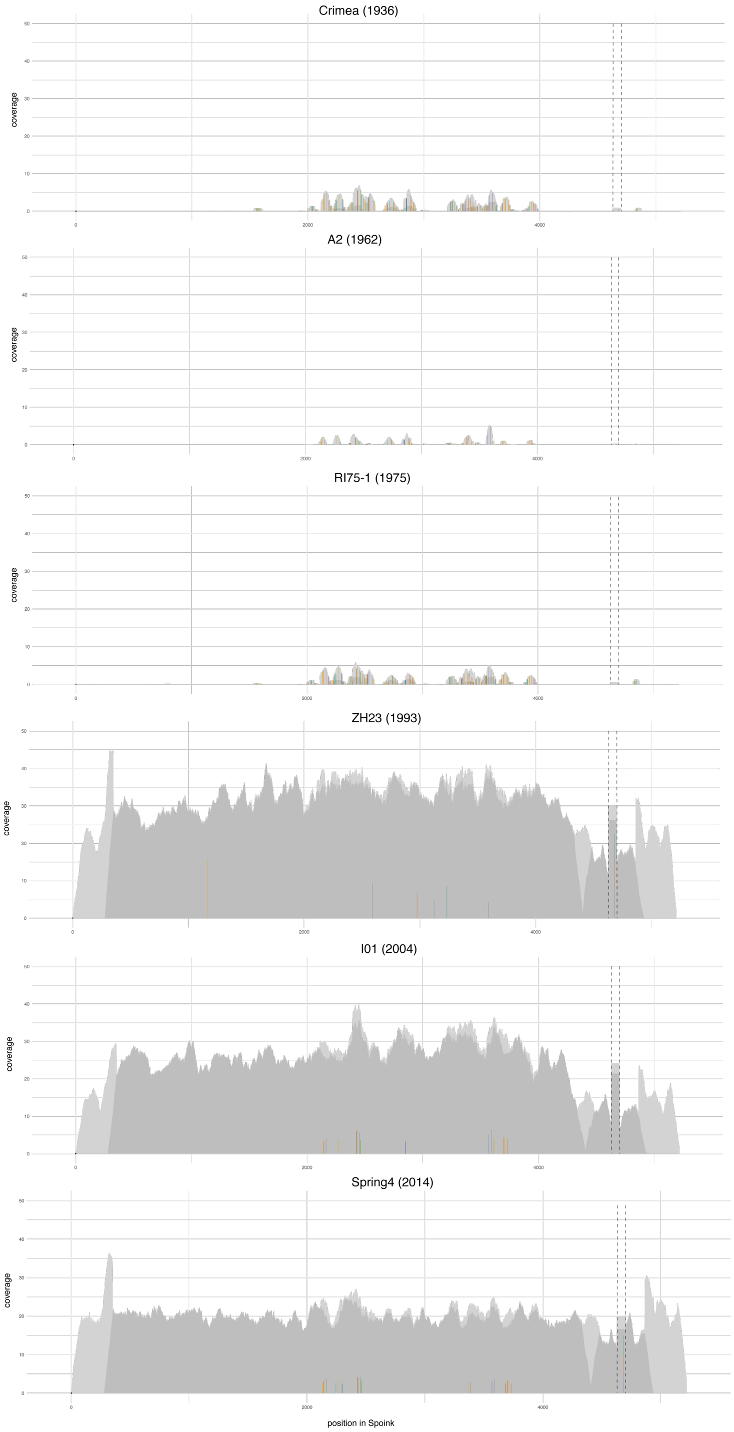


Figure 2: DeviaTE plots of six *D. melanogaster* strains collected during the last century. The short reads were aligned to the consensus sequence of *Spoink* and the coverage was normalized to the the coverage of single-copy genes. The coverage was manually curbed at the poly-A track (indicated by dashed lines). Note that very few reads of old strains ($\leq$ 1975) align to *Spoink* whereas a contiguous coverage of reads along *Spoink* is observed for more recently collected strains ($\geq$ 1993).

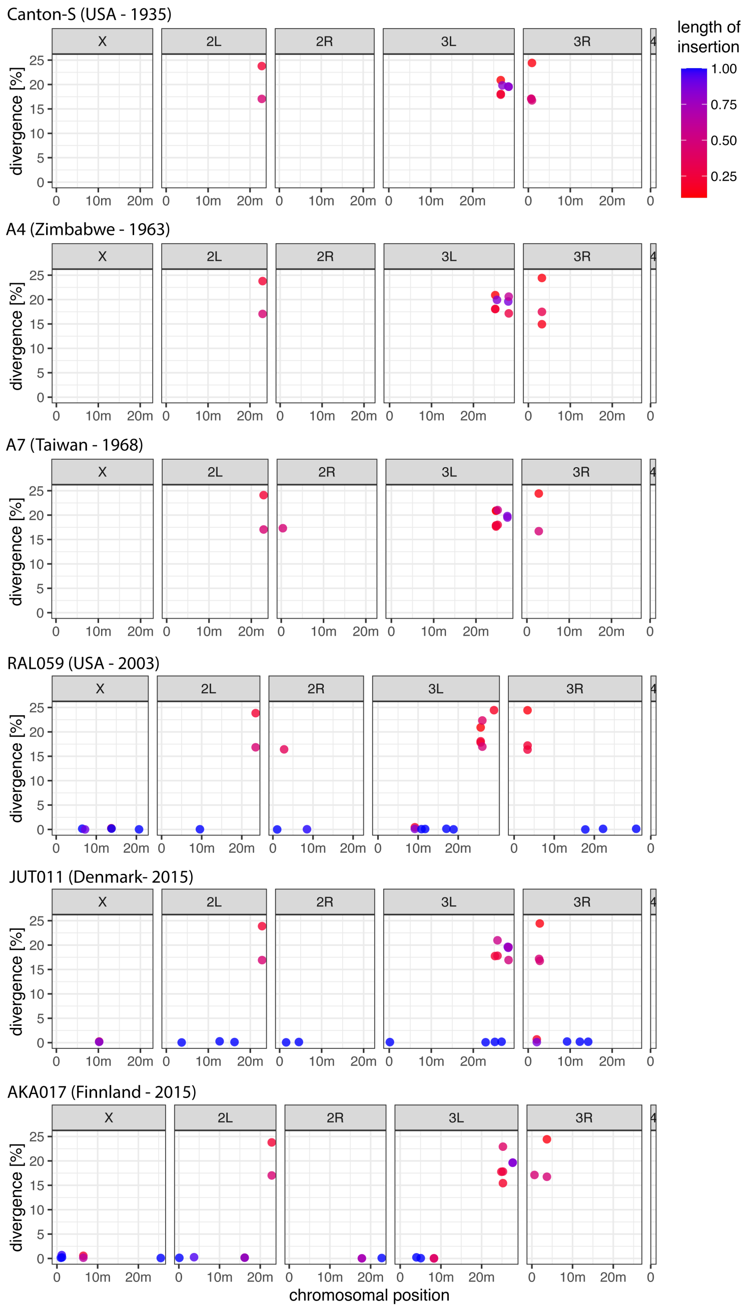


Figure 3: Abundance of *Spoink* insertions in six long-read assemblies of *D. melanogaster* strains collected during the last century. Note that all strains contain fragmented and diverged insertions of *Spoink*, while solely recently collected strains ($\geq$ 2003) contain canonical *Spoink* insertions (i.e. full-length insertions with little divergence from the consensus sequence). .

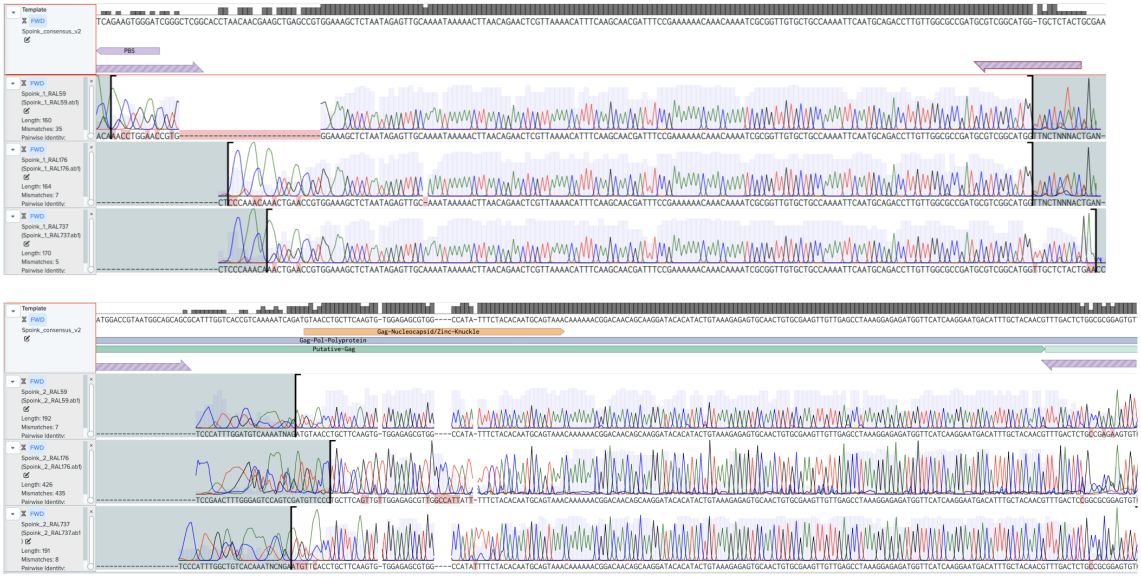


Figure 4: The Sanger sequence of the six PCR amplicons matches with the consensus sequence of Spoink. The Sanger sequences of the amplicons amplified by P1 (top) and P2 (bottom) were aligned to the sequence of *Spoink* using Benchling. Three amplicons were sequenced for each primer pair. The position of the PCR primers are shown at the top of each panel (purple arrows).

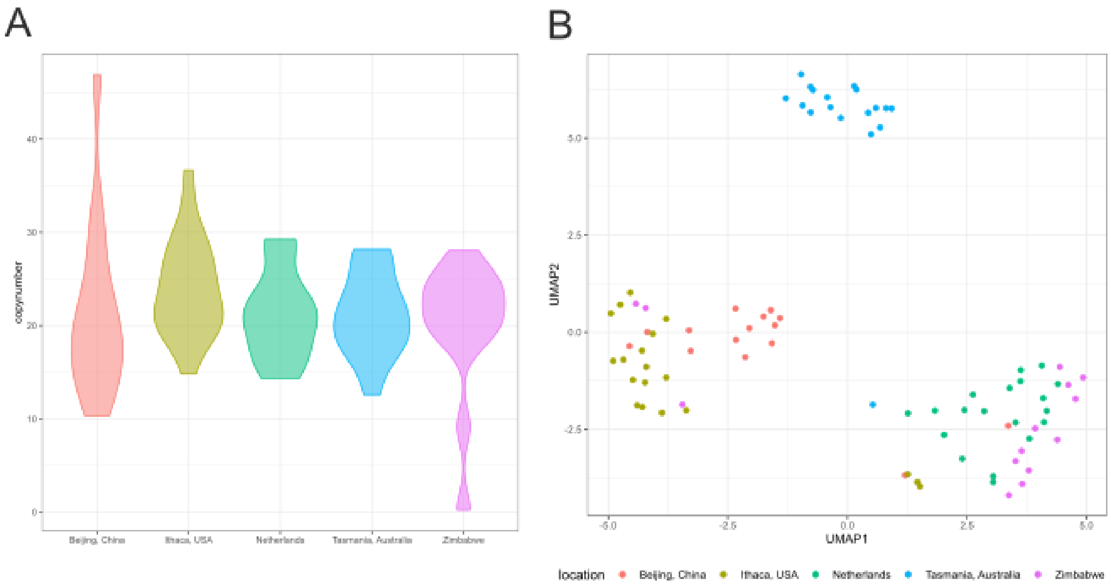


Figure 5: Abundance and composition of *Spoink* insertions in the GDL. A) Abundance of *Spoink* in the GDL. Note that one strain from Zimbabwe does not have any *Spoink* insertion. B) UMAP summarizing the composition of *Spoink* among the GDL. Note that *Spoink* shows a pronounced population structure, where three main clusters can be discerned: Tasmania, Bejing/Ithaca and Netherlands/Zimbabwe.

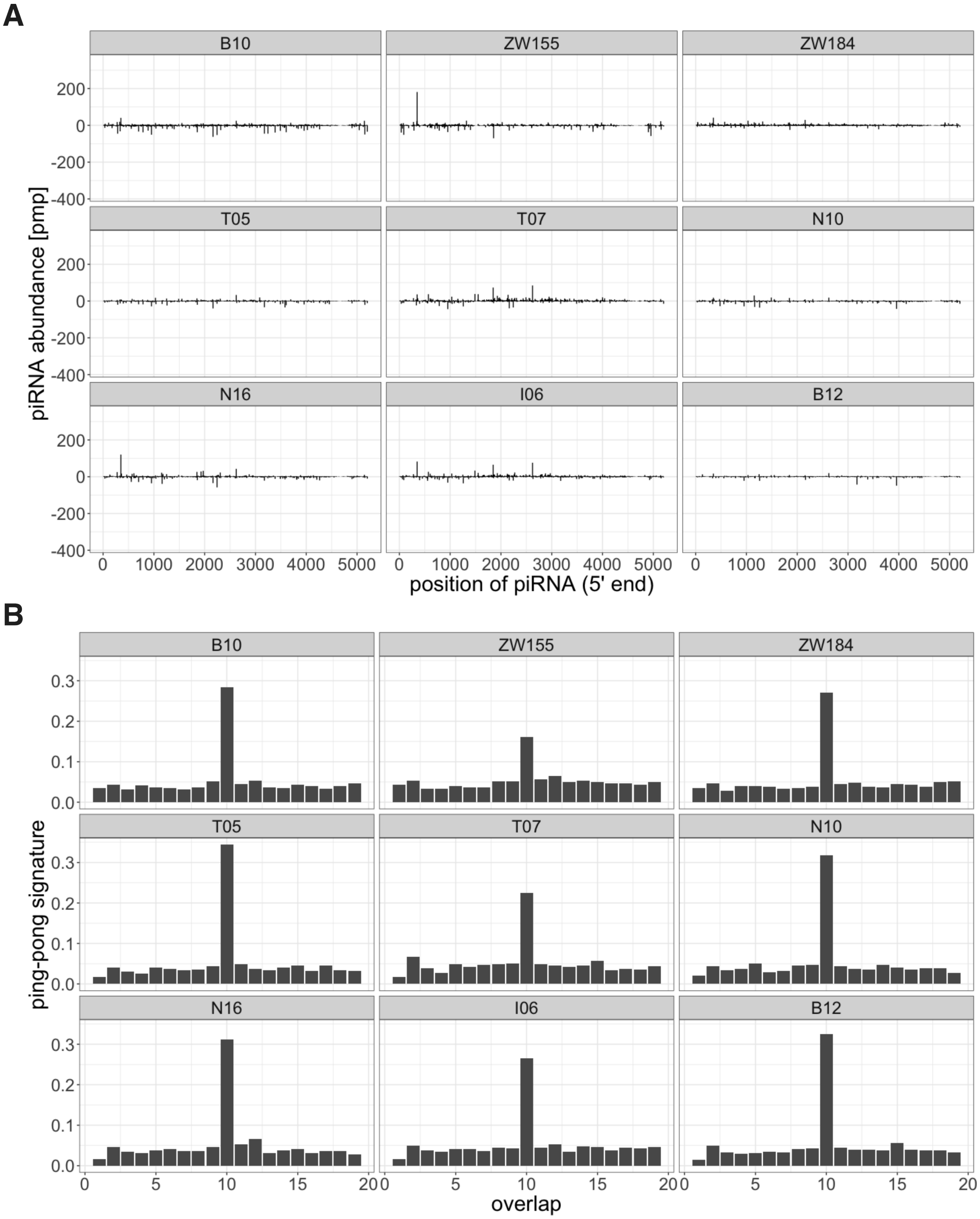


Figure 6: A piRNA based defence against *Spoink* is active in the 10 GDL strains. Two strains are analysed for each continent (Bxx Beijing/Asia, Ixx Ithaca/America, Nxx Netherlands/Europe, Txx Tasmania/Australia, ZWxx Zimbabwe/Africa; the second strain from Ithaca (*I17*) is shown in the main manuscript). A) piRNAs mapping to the sequence of *Spoink*. Solely the 5' positions of piRNAs are shown and the piRNA abundance is normalized to one million piRNAs. Sense piRNAs are shown on the positive y-axis and antisense piRNAs on the negative y-axis. B) ping-pong signature of *Spoink*.

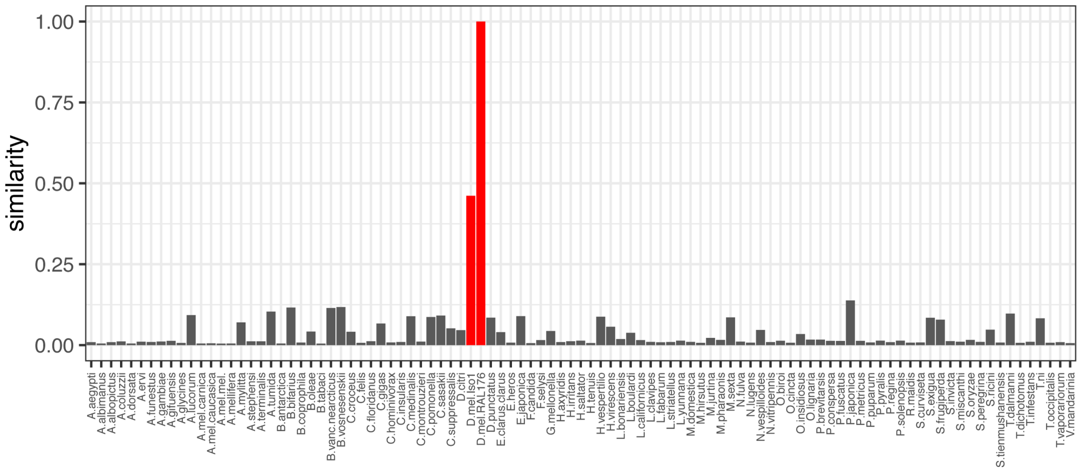


Figure 7: Barplots show the similarity between the consensus sequence of *Spoink* and the best match in each of 99 long-read assemblies of diverse insect species. As a reference, two *D. melanogaster* assemblies (red) were included, where D.mel.RAL176 has canonical *Spoink* insertions while D.mel.Iso1 solely has degraded fragments of sequences having some similarity with *Spoink*.

#### Supplementary tables

Table 1: We discovered *Spoink* by noticing differences in the abundance of *Gypsy_7_DEl* between the reference genome Iso1 and a long-read assemblies from a more recently collected strain (TOM007 was collected in 2015 [Rech et al., 2022]). The best ten matches for *Gypsy_7_DEl* and the consensus sequence of *Spoink* are shown for both assemblies. Matches were identified with RepeatMasker [Smit et al., 1996-2010]. Note that the discrepancy between Iso1 and TOM007 is more pronounced when the consensus sequence of *Spoink* is considered.

| score | subst | len | chrm | start | end | rc | TE | assembly |
|---|---|---|---|---|---|---|---|---|
| 23487 | 16.32 | 0.9 | JAEIGS010000237.1 | 513885 | 517979 | + | Gypsy_7_DEl | Iso1 |
| 21209 | 16.57 | 0.88 | JAEIGS010000286.1 | 947301 | 951265 | + | Gypsy_7_DEl | Iso1 |
| 15961 | 17.56 | 0.61 | JAEIGS010000237.1 | 590996 | 593798 | + | Gypsy_7_DEl | Iso1 |
| 6055 | 20.06 | 0.26 | JAEIGS010000089.1 | 299475 | 300630 | + | Gypsy_7_DEl | Iso1 |
| 4233 | 19.69 | 0.18 | JAEIGS010000051.1 | 1106 | 1938 | + | Gypsy_7_DEl | Iso1 |
| 3665 | 18.84 | 0.16 | JAEIGS010000286.1 | 940239 | 940967 | + | Gypsy_7_DEl | Iso1 |
| 2230 | 19.37 | 0.09 | JAEIGS010000110.1 | 46405 | 46817 | + | Gypsy_7_DEl | Iso1 |
| 1073 | 16.48 | 0.09 | JAEIGS010000254.1 | 13281 | 13644 | + | Gypsy_7_DEl | Iso1 |
| 1054 | 16.99 | 0.09 | JAEIGS010000110.1 | 49397 | 49755 | + | Gypsy_7_DEl | Iso1 |
| 925 | 18.75 | 0.04 | JAEIGS010000286.1 | 539354 | 539529 | C | Gypsy_7_DEl | Iso1 |
| 24069 | 16.8 | 0.9 | CM034738.1 | 394890 | 399008 | + | Gypsy_7_DEl | TOM007 |
| 23424 | 18.89 | 0.92 | CM034732.1 | 20134893 | 20139139 | C | Gypsy_7_DEl | TOM007 |
| 23408 | 16.29 | 0.9 | CM034735.1 | 28195698 | 28199789 | + | Gypsy_7_DEl | TOM007 |
| 23254 | 18.83 | 0.91 | CM034735.1 | 10927462 | 10931666 | C | Gypsy_7_DEl | TOM007 |
| 23215 | 18.89 | 0.92 | CM034732.1 | 20106718 | 20110964 | C | Gypsy_7_DEl | TOM007 |
| 23174 | 18.9 | 0.9 | CM034732.1 | 15164322 | 15168487 | C | Gypsy_7_DEl | TOM007 |
| 23165 | 18.85 | 0.9 | CM034735.1 | 6916392 | 6920556 | C | Gypsy_7_DEl | TOM007 |
| 23128 | 19 | 0.9 | CM034734.1 | 21517228 | 21521397 | C | Gypsy_7_DEl | TOM007 |
| 23074 | 19.02 | 0.9 | CM034733.1 | 2859281 | 2863449 | + | Gypsy_7_DEl | TOM007 |
| 23074 | 19.02 | 0.9 | CM034733.1 | 6071040 | 6075208 | + | Gypsy_7_DEl | TOM007 |
| 22322 | 19.61 | 0.87 | JAEIGS010000237.1 | 513636 | 518077 | + | Spoink | Iso1 |
| 19827 | 19.94 | 0.83 | JAEIGS010000286.1 | 947069 | 951265 | + | Spoink | Iso1 |
| 14846 | 20.61 | 0.59 | JAEIGS010000237.1 | 590775 | 593762 | + | Spoink | Iso1 |
| 14188 | 16.75 | 0.47 | JAEIGS010000110.1 | 47013 | 49449 | + | Spoink | Iso1 |
| 13357 | 16.91 | 0.46 | JAEIGS010000089.1 | 300606 | 302918 | + | Spoink | Iso1 |
| 13151 | 17.2 | 0.47 | JAEIGS010000254.1 | 10796 | 13234 | + | Spoink | Iso1 |
| 9332 | 28.9 | 0.6 | JAEIGS010000104.1 | 400756 | 403829 | + | Spoink | Iso1 |
| 8667 | 33.23 | 0.76 | JAEIGS010000117.1 | 184757 | 188686 | + | Spoink | Iso1 |
| 8098 | 32.34 | 0.76 | JAEIGS010000049.1 | 24780858 | 24784790 | + | Spoink | Iso1 |
| 7762 | 31.6 | 0.7 | JAEIGS010000059.1 | 470089 | 473701 | + | Spoink | Iso1 |
| 48688 | 0 | 1 | CM034735.1 | 6915682 | 6920905 | C | Spoink | TOM007 |
| 48616 | 0.02 | 1 | CM034734.1 | 21308387 | 21313610 | + | Spoink | TOM007 |
| 48613 | 0.02 | 1 | CM034732.1 | 15163600 | 15168836 | C | Spoink | TOM007 |
| 48573 | 0.06 | 1 | CM034733.1 | 6070691 | 6075920 | + | Spoink | TOM007 |
| 48507 | 0.1 | 1 | CM034733.1 | 19565354 | 19570589 | C | Spoink | TOM007 |
| 48498 | 0.04 | 1 | CM034733.1 | 2858932 | 2864175 | + | Spoink | TOM007 |
| 48434 | 0 | 1 | CM034735.1 | 21881386 | 21886609 | + | Spoink | TOM007 |
| 48410 | 0 | 1 | CM034734.1 | 21516518 | 21521746 | C | Spoink | TOM007 |
| 48123 | 0.13 | 1 | CM034735.1 | 10926746 | 10932015 | C | Spoink | TOM007 |
| 46927 | 0.09 | 1 | CM034734.1 | 1659119 | 1664406 | C | Spoink | TOM007 |

Table 2: Similarity between *Spoink* and other TEs in the different repeat libraries generated for *D. melanogaster*. For each repeat library the best five hits are shown. Solely matches with a minimum overlap with *Spoink* of at least 30% are considered. subst. substitions in percent between *Spoink* and the TE, len. fraction of the length of a TE aligning with *Spoink*; a Rech et al. [2022], b Quesneville et al. [2005], c Chakraborty et al. [2021], d Srivastav et al. [2023], e Ellison and Cao [2020]

| subst. | len. | TE | source |
|---|---|---|---|
| 0.13 | 0.87 | con41_UnFmcl001_RLX-incomp | a |
| 19.34 | 0.85 | con4_UnFmcl034_RLX-incomp | a |
| 32.22 | 0.76 | con8_micropia | a |
| 31.04 | 0.59 | con13_UnFmcl013_RLX-incomp | a |
| 32.08 | 0.48 | con4_invader6 | a |
| 32.52 | 0.76 | DMDM11 (Micropia) | b |
| 31.34 | 0.33 | INVADER6 | b |
| 18.87 | 0.80 | Gypsy_7_DEl_I_LTR_Gypsy | c |
| 17.93 | 0.67 | Gypsy_2_DSim_I_LTR_Gypsy | c |
| 29.4 | 0.78 | Gypsy9_I_Dya_LTR_Gypsy | c |
| 30.26 | 0.82 | Gypsy_38_DAn_I_LTR_Gypsy | c |
| 30.25 | 0.79 | Gypsy_49_DWil_I_LTR_Gypsy | c |
| 29.97 | 0.54 | family-58_micropia-like | d |
| 32.52 | 0.76 | micropia | d |
| 30.1 | 0.57 | family-63_micropia-like | d |
| 31.42 | 0.61 | MN418888.1 | e |

Table 3: Identity of sequences in Oregon-R having some similarity with the consensus sequence of *Spoink*. Solely sequences having a divergence of $\leq$ 25% and minimum overlap of at least 10% with *Spoink* are considered. The sequences were extracted from the assembly of Oregon-R (chromosome:start-end) and aligned against the TE library of *D. melanogaster* using blastn [Altschul et al., 1990, Quesneville et al., 2005]. Most of these sequences match TARTC and DMDM11.

| chromosome:start-end | TE | len. | identity |
|---|---|---|---|
| CM010514.1:22763163-22771454 | DMDM11(Micropia) | 25% | 64% |
| CM010516.1:24739316-24746262 | DME487856 (Max-element) | 26% | 92% |
| CM010516.1:25103425-25110397 | TARTC | 25% | 61% |
| CM010516.1:25110536-25117413 | TARTC | 25% | 71% |
| CM010516.1:25129800-25137879 | DMDM11(Micropia) | 33% | 65% |
| CM010516.1:25132367-25140446 | DMDM11(Micropia) | 56% | 65% |
| CM010516.1:25134448-25142814 | DMDM11(Micropia) | 47% | 65% |
| CM010516.1:27328230-27337103 | DMDM11(Micropia) | 39% | 65% |
| CM010516.1:27329110-27338574 | DMDM11(Micropia) | 37% | 65% |
| CM010517.1:2455875-2462520 | DMDM11(Micropia) | 38% | 67% |
| CM010517.1:2456731-2465167 | DMDM11(Micropia) | 31% | 67% |
| PGRW01000073.1:0-5565 | DMDM11(Micropia) | 39% | 67% |
| PGRW01000073.1:0-5565 | DMDM11(Micropia) | 39% | 67% |

Table 4: Overview of the short-read data analysed in this work. Data are from Grenier et al. [2015], Schwarz et al. [2021], Long et al. [2013], Lange et al. [2021], Rech et al. [2022])

| accession | strain | year | location |
|---|---|---|---|
| SRR23876563 | museum | 1800 | Lund, Sweden |
| SRR23876564 | museum | 1800 | Lund, Sweden |
| SRR23876562 | museum | 1850 | Passau, Germany |
| SRR23876569 | museum | 1850 | Passau, Germany |
| SRR23876586 | museum | 1933 | Lund, Sweden |
| SRR11846555 | Crimea | 1936 | Crimea, Ukraine |
| SRR11846565 | Hikone-R | 1958 | Japan |
| SRR457698 | A2(BOG1) | 1962 | Bogota, Colombia |
| SRR457707 | A4(KSA2) | 1963 | South Africa |
| SRR457701 | B4(RVC3) | 1963 | California, USA |
| SRR457669 | A5(VAG1) | 1965 | Athens, Greece |
| SRR457697 | A6(wild5B) | 1966 | Georgia, USA |
| SRR11846560 | Harwich | 1967 | Harwich, MA, USA |
| SRR11460801 | Pi2 | 1975 | |
| SRR13257684 | RI75-1 | 1975 | Providence, USA |
| SRR14293191 | RI75-2 | 1975 | Providence, USA |
| SRR14293574 | RI75-5 | 1975 | Providence, USA |
| SRR14293140 | RI75-10 | 1975 | Providence, USA |
| SRR14293233 | RI75-4 | 1975 | Providence, USA |
| SRR14294393 | RI75-7 | 1975 | Providence, USA |
| SRR14294796 | RI75-11 | 1975 | Providence, USA |
| SRR14294940 | RI75-13 | 1975 | Providence, USA |
| SRR14293576 | RI75-6 | 1975 | Providence, USA |
| SRR14294899 | RI75-12 | 1975 | Providence, USA |
| SRR14294944 | RI77-50 | 1977 | Providence, USA |
| SRR14296646 | RI77-52 | 1977 | Providence, USA |
| SRR14296993 | RI77-53 | 1977 | Providence, USA |
| SRR14297433 | RI77-61 | 1977 | Providence, USA |
| SRR14297455 | RI77-55 | 1977 | Providence, USA |
| SRR14298094 | RI77-60 | 1977 | Providence, USA |
| SRR14296430 | RI7751 | 1977 | Providence, USA |
| SRR14297771 | RI77-58 | 1977 | Providence, USA |
| SRR14297795 | RI77-56 | 1977 | Providence, USA |
| SRR14298064 | RI77-54 | 1977 | Providence, USA |
| SRR14297441 | RI78-1A | 1978 | Providence, USA |
| SRR14297770 | RI78-6 | 1978 | Providence, USA |
| SRR14297954 | RI78-14 | 1978 | Providence, USA |
| SRR14298028 | RI78-1B | 1978 | Providence, USA |
| SRR14297434 | RI78-12 | 1978 | Providence, USA |
| SRR14297796 | RI78-11 | 1978 | Providence, USA |
| SRR14297813 | RI78-8 | 1978 | Providence, USA |
| SRR14297794 | RI78-13 | 1978 | Providence, USA |
| SRR14298002 | RI78-5 | 1978 | Providence, USA |
| SRR14298055 | RI78-7 | 1978 | Providence, USA |
| SRR14298091 | RI78-9 | 1978 | Providence, USA |
| SRR14297456 | RI79-7 | 1979 | Providence, USA |
| SRR14298093 | RI79-3 | 1979 | Providence, USA |
| SRR14306823 | RI79-10 | 1979 | Providence, USA |
| SRR14306826 | RI79-17 | 1979 | Providence, USA |
| SRR14306827 | RI79-16 | 1979 | Providence, USA |
| SRR14306825 | RI79-19 | 1979 | Providence, USA |
| SRR14306829 | RI79-14 | 1979 | Providence, USA |
| SRR14306830 | RI79-13 | 1979 | Providence, USA |
| SRR14298105 | RI79-1 | 1979 | Providence, USA |
| SRR14306822 | RI79-20 | 1979 | Providence, USA |
| SRR14306824 | RI79-9 | 1979 | Providence, USA |
| SRR14306828 | RI79-15 | 1979 | Providence, USA |
| SRR14306831 | RI79-12 | 1979 | Providence, USA |
| SRR14306842 | RI79-11 | 1979 | Providence, USA |
| SRR14306841 | RI80-15 | 1980 | Providence, USA |
| SRR14306844 | RI80-11 | 1980 | Providence, USA |
| SRR14306847 | RI80-7 | 1980 | Providence, USA |
| SRR14306850 | RI80-4 | 1980 | Providence, USA |
| SRR14306821 | RI80-2 | 1980 | Providence, USA |
| SRR14306843 | RI80-13 | 1980 | Providence, USA |
| SRR14306845 | RI80-9 | 1980 | Providence, USA |
| SRR14306846 | RI80-8 | 1980 | Providence, USA |
| SRR14306848 | RI80-6 | 1980 | Providence, USA |
| SRR14306849 | RI80-5 | 1980 | Providence, USA |
| SRR14306833 | RI83-8 | 1983 | Providence, USA |
| SRR14306835 | RI83-6 | 1983 | Providence, USA |
| SRR14306837 | RI83-4 | 1983 | Providence, USA |
| SRR14306838 | RI83-3 | 1983 | Providence, USA |
| SRR14306834 | RI83-7 | 1983 | Providence, USA |
| SRR14306832 | RI83-9 | 1983 | Providence, USA |
| SRR14306836 | RI83-5 | 1983 | Providence, USA |
| SRR14306839 | RI83-2 | 1983 | Providence, USA |
| SRR14306840 | RI83-1 | 1983 | Providence, USA |
| SRR1663598 | ZH23 | 1993 | Zimbabwe |
| SRR1663599 | ZH26 | 1993 | Zimbabwe |
| SRR1663600 | ZH33 | 1993 | Zimbabwe |
| SRR1663601 | ZH42 | 1993 | Zimbabwe |
| SRR1663602 | ZS10 | 1993 | Zimbabwe |
| SRR1663603 | ZW09 | 1993 | Zimbabwe |
| SRR1663604 | ZW139 | 1993 | Zimbabwe |
| SRR1663605 | ZW140 | 1993 | Zimbabwe |
| SRR1663606 | ZW142 | 1993 | Zimbabwe |
| SRR1663607 | ZW144 | 1993 | Zimbabwe |
| SRR1663608 | ZW155 | 1993 | Zimbabwe |
| SRR1663609 | ZW177 | 1993 | Zimbabwe |
| SRR1663610 | ZW184 | 1993 | Zimbabwe |
| SRR1663611 | ZW185 | 1993 | Zimbabwe |
| SRR1769729 | ZW155 | 1993 | Zimbabwe |
| SRR1662283 | B04 | 1995 | Beijing, China |
| SRR1663528 | B05 | 1995 | Beijing, China |
| SRR1663529 | B10 | 1995 | Beijing, China |
| SRR1663530 | B11 | 1995 | Beijing, China |
| SRR1663531 | B12 | 1995 | Beijing, China |
| SRR1663532 | B14 | 1995 | Beijing, China |
| SRR1663533 | B23 | 1995 | Beijing, China |
| SRR1663534 | B28 | 1995 | Beijing, China |
| SRR1663535 | B38 | 1995 | Beijing, China |
| SRR1663536 | B42 | 1995 | Beijing, China |
| SRR1663537 | B43 | 1995 | Beijing, China |
| SRR1663538 | B51 | 1995 | Beijing, China |
| SRR1663539 | B52 | 1995 | Beijing, China |
| SRR1663540 | B54 | 1995 | Beijing, China |
| SRR1663541 | B59 | 1995 | Beijing, China |
| SRR1663561 | N01 | 2003 | Netherlands |
| SRR1663562 | N02 | 2003 | Netherlands |
| SRR1663563 | N03 | 2003 | Netherlands |
| SRR1663564 | N04 | 2003 | Netherlands |
| SRR1663565 | N07 | 2003 | Netherlands |
| SRR1663566 | N10 | 2003 | Netherlands |
| SRR1663567 | N11 | 2003 | Netherlands |
| SRR1663568 | N13 | 2003 | Netherlands |
| SRR1663569 | N14 | 2003 | Netherlands |
| SRR1663570 | N15 | 2003 | Netherlands |
| SRR1663571 | N16 | 2003 | Netherlands |
| SRR1663572 | N17 | 2003 | Netherlands |
| SRR1663573 | N18 | 2003 | Netherlands |
| SRR1663574 | N19 | 2003 | Netherlands |
| SRR1663575 | N22 | 2003 | Netherlands |
| SRR1663576 | N23 | 2003 | Netherlands |
| SRR1663577 | N25 | 2003 | Netherlands |
| SRR1663578 | N29 | 2003 | Netherlands |
| SRR1663579 | N30 | 2003 | Netherlands |
| SRR1663580 | T01 | 2003 | Tasmania, Australia |
| SRR1663581 | T04 | 2003 | Tasmania, Australia |
| SRR1663582 | T05 | 2003 | Tasmania, Australia |
| SRR1663583 | T07 | 2003 | Tasmania, Australia |
| SRR1663584 | T09 | 2003 | Tasmania, Australia |
| SRR1663585 | T10 female | 2003 | Tasmania, Australia |
| SRR1663586 | T14A | 2003 | Tasmania, Australia |
| SRR1663587 | T22A | 2003 | Tasmania, Australia |
| SRR1663588 | T23 | 2003 | Tasmania, Australia |
| SRR1663589 | T24 | 2003 | Tasmania, Australia |
| SRR1663590 | T25A | 2003 | Tasmania, Australia |
| SRR1663591 | T29A | 2003 | Tasmania, Australia |
| SRR1663592 | T30 | 2003 | Tasmania, Australia |
| SRR1663593 | T35 | 2003 | Tasmania, Australia |
| SRR1663594 | T36B | 2003 | Tasmania, Australia |
| SRR1663595 | T39 | 2003 | Tasmania, Australia |
| SRR1663596 | T43A | 2003 | Tasmania, Australia |
| SRR1663597 | T45B | 2003 | Tasmania, Australia |
| SRR1663542 | I01 | 2004 | Ithaca, USA |
| SRR1663543 | I02 | 2004 | Ithaca, USA |
| SRR1663544 | I03 | 2004 | Ithaca, USA |
| SRR1663545 | I04 | 2004 | Ithaca, USA |
| SRR1663546 | I06 | 2004 | Ithaca, USA |
| SRR1663547 | I07 | 2004 | Ithaca, USA |
| SRR1663548 | I13 | 2004 | Ithaca, USA |
| SRR1663549 | I16 | 2004 | Ithaca, USA |
| SRR1663550 | I17 | 2004 | Ithaca, USA |
| SRR1663551 | I22 | 2004 | Ithaca, USA |
| SRR1663552 | I23 | 2004 | Ithaca, USA |
| SRR1663553 | I24 | 2004 | Ithaca, USA |
| SRR1663554 | I26 | 2004 | Ithaca, USA |
| SRR1663555 | I29 | 2004 | Ithaca, USA |
| SRR1663556 | I31 | 2004 | Ithaca, USA |
| SRR1663557 | I33 | 2004 | Ithaca, USA |
| SRR1663558 | I34 | 2004 | Ithaca, USA |
| SRR1663559 | I35 | 2004 | Ithaca, USA |
| SRR1663560 | I38 | 2004 | Ithaca, USA |
| SRR14308759 | spring4 | 2014 | Providence, USA |
| SRR14308763 | fall6 | 2014 | Providence, USA |
| SRR14308767 | spring12 | 2014 | Providence, USA |
| SRR14308768 | spring11 | 2014 | Providence, USA |
| SRR14308772 | spring7 | 2014 | Providence, USA |
| SRR14308776 | fall1 | 2014 | Providence, USA |
| SRR14308762 | spring1 | 2014 | Providence, USA |
| SRR14308764 | fall5 | 2014 | Providence, USA |
| SRR14308766 | fall3 | 2014 | Providence, USA |
| SRR14308774 | spring5 | 2014 | Providence, USA |
| SRR14308775 | fall2 | 2014 | Providence, USA |
| SRR14308760 | spring3 | 2014 | Providence, USA |
| SRR14308761 | spring2 | 2014 | Providence, USA |
| SRR14308765 | fall4 | 2014 | Providence, USA |
| SRR14308769 | spring10 | 2014 | Providence, USA |
| SRR14308770 | spring9 | 2014 | Providence, USA |
| SRR14308771 | spring8 | 2014 | Providence, USA |
| SRR14308773 | spring6 | 2014 | Providence, USA |
| SRR9951090 | ES_Ten_15_15 | 2015 | Tenerife, Spain |

Table 5: Overview of the long-read assemblies of *D. melanogaster* strains analysed in this work. For each strain we show the assembly ID, the strain, the sampling location and the sampling date. a King et al. [2012], Chakraborty et al. [2019], b Wierzbicki et al. [2021], c Hoskins et al. [2015], d Rech et al. [2022], e Mackay et al. [2012]

| assembly | strain | location | time | source |
|---|---|---|---|---|
| GCA_003397115.2 | B6 | Ica/Peru | 1956 | a |
| GCA_003401685.1 | AB8 | na | na | a |
| GCA_003401735.1 | Canton-S | Ohio/USA | 1935 | a |
| GCA_003401745.1 | A4 | Koriba Dam/Zimbabwe | 1963 | a |
| GCA_003401795.1 | A3 | Barcelona/Spain | 1954 | a |
| GCA_003401805.1 | A2 | Bogota/Columbia | 1962 | a |
| GCA_003401855.1 | A5 | Athens/Greece | 1965 | a |
| GCA_003401885.1 | A6 | Georgia/USA | 1966 | a |
| GCA_003401915.1 | A7 | Ken-ting/Taiwan | 1968 | a |
| GCA_003401925.1 | B1 | Bermuda | 1954 | a |
| GCA_003401975.1 | B2 | Capetown/SA | 1954 | a |
| GCA_003402005.1 | B4 | California/USA | 1963 | a |
| GCA_003402015.1 | Oregon-R | Oregon/USA | 1925 | a |
| GCA_003402055.1 | B3 | Israel | 1954 | a |
| GCA_015852585.1 | Pi2 | na | 1975 | b |
| GCA_000001215.4 | Iso1 | na | na | c |
| GCA_015832445.1 | Canton-S | Ohio/USA | 1935 | b |
| GCA_020142105.1 | AKA017 | Akka/Finland | 2015 | d |
| GCA_020169495.1 | AKA018 | Akka/Finland | 2015 | d |
| GCA_020142085.1 | COR014 | Cortes de Baza/Spain | 2015 | d |
| GCA_020142045.1 | COR018 | Cortes de Baza/Spain | 2015 | d |
| GCA_020142055.1 | COR023 | Cortes de Baza/Spain | 2015 | d |
| GCA_020142025.1 | COR025 | Cortes de Baza/Spain | 2015 | d |
| GCA_020141985.1 | GIM012 | Gimenells/Spain | 2015 | d |
| GCA_020142005.1 | GIM024 | Gimenells/Spain | 2015 | d |
| GCA_020141955.1 | JUT008 | Jutland/Denmark | 2015 | d |
| GCA_020141935.1 | JUT011 | Jutland/Denmark | 2015 | d |
| GCA_020141925.1 | KIE094 | Kiev/Ucraina | 2018 | d |
| GCA_020141875.1 | LUN004 | Lund/Sweden | 2015 | d |
| GCA_020141855.1 | LUN007 | Lund/Sweden | 2015 | d |
| GCA_020141845.1 | MUN008 | Munich/Germany | 2015 | d |
| GCA_020141835.1 | MUN009 | Munich/Germany | 2015 | d |
| GCA_020141815.1 | MUN013 | Munich/Germany | 2015 | d |
| GCA_020141795.1 | MUN015 | Munich/Germany | 2015 | d |
| GCA_020141765.1 | MUN016 | Munich/Germany | 2015 | d |
| GCA_020141735.1 | MUN020 | Munich/Germany | 2015 | d |
| GCA_020141745.1 | RAL059 | Raleigh/USA | 2003 | d,e |
| GCA_020141705.1 | RAL091 | Raleigh/USA | 2003 | d,e |
| GCA_020141665.1 | RAL176 | Raleigh/USA | 2003 | d,e |
| GCA_020141655.1 | RAL177 | Raleigh/USA | 2003 | d,e |
| GCA_020141675.1 | RAL375 | Raleigh/USA | 2003 | d,e |
| GCA_020141625.1 | RAL426 | Raleigh/USA | 2003 | d,e |
| GCA_020141585.1 | RAL737 | Raleigh/USA | 2003 | d,e |
| GCA_020141575.1 | RAL855 | Raleigh/USA | 2003 | d,e |
| GCA_020141595.1 | SLA001 | Slankamen/Serbia | 2016 | d |
| GCA_020141495.1 | STO022 | Stockholm/Sweden | 2011 | d |
| GCA_020141505.1 | TEN015 | Tenerife/Spain | 2015 | d |
| GCA_020141485.1 | TOM007 | Tomelloso/Spain | 2015 | d |
| GCA_020141515.1 | TOM008 | Tomelloso/Spain | 2015 | d |

Table 6: *Spoink* insertions in piRNA clusters of long-read assemblies of different *D. melanogaster* strains [Rech et al., 2022]. Note that for several strains we could not find a single *Spoink* insertion in a piRNA cluster. On the other hand, some strains, like RAL176, have multiple *Spoink* insertions in piRNA clusters.

| | Spoink | | | piRNA cluster | | | | |
|---|---|---|---|---|---|---|---|---|
| Strain | chr | start | end | chr | start | end | ID | Overlap |
| AK017 | NA | NA | NA | NA | NA | NA | NA | NA |
| AK018 | NA | NA | NA | NA | NA | NA | NA | NA |
| COR014 | NA | NA | NA | NA | NA | NA | NA | NA |
| COR018 | CM034904.1 | 1,451,740 | 1,456,874 | CM034904.1 | 1,433,545 | 1,478,569 | 137 | 5,134 |
| COR023 | CM034914.1 | 22,928,374 | 22,933,154 | CM034914.1 | 22,891,335 | 22,953,549 | 45 | 4,780 |
| COR023 | CM034914.1 | 22,933,255 | 22,937,256 | CM034914.1 | 22,891,335 | 22,953,549 | 45 | 4,001 |
| COR025 | NA | NA | NA | NA | NA | NA | NA | NA |
| RAL426 | CM034769.1 | 7,091,652 | 7,096,769 | CM034769.1 | 7,001,338 | 7,299,749 | 1 | 5,117 |
| GIM012 | NA | NA | NA | NA | NA | NA | NA | NA |
| GIM024 | NA | NA | NA | NA | NA | NA | NA | NA |
| JUT00008 | CM034879.1 | 24,998,160 | 25,003,388 | CM034879.1 | 24,972,757 | 25,022,954 | 45 | 5,228 |
| JU011 | CM034875.1 | 26,745,760 | 26,751,014 | CM034875.1 | 26,739,413 | 26,932,771 | 12 | 5,254 |
| JU011 | CM034875.1 | 25,143,543 | 25,148,792 | CM034875.1 | 25,088,014 | 25,187,432 | 15 | 5,249 |
| JU011 | CM034875.1 | 25,143,543 | 25,148,792 | CM034875.1 | 25,061,149 | 26,113,760 | 24 | 5,249 |
| KIE094 | NA | NA | NA | NA | NA | NA | NA | NA |
| LUN0004 | NA | NA | NA | NA | NA | NA | NA | NA |
| LUN007 | NA | NA | NA | NA | NA | NA | NA | NA |
| MUN008 | CM034853.1 | 5,026,490 | 5,031,753 | CM034853.1 | 4,917,190 | 5,062,654 | 28 | 5,263 |
| MUN008 | CM034851.1 | 22,521,456 | 22,526,701 | CM034851.1 | 21,902,471 | 22,858,985 | 8_9 | 5,245 |
| MUN009 | NA | NA | NA | NA | NA | NA | NA | NA |
| MUN003 | CM034840.1 | 12,763,342 | 12,768,634 | CM034840.1 | 10,535,909 | 25,146,430 | 15 | 5,292 |
| MUN015 | CM034825.1 | 6,963,416 | 6,968,674 | CM034825.1 | 6,856,930 | 7,159,555 | 1 | 5,258 |
| MUN016 | NA | NA | NA | NA | NA | NA | NA | NA |
| MUN020 | CM034813.1 | 2,611,417 | 2,616,648 | CM034813.1 | 2,021,000 | 3,141,406 | 30 | 5,231 |
| MUN020 | CM034813.1 | 2,611,417 | 2,616,648 | CM034813.1 | 2,058,138 | 3,079,295 | 93 | 5,231 |
| MUN020 | CM034812.1 | 25,689,940 | 25,695,151 | CM034812.1 | 25,390,627 | 26,359,124 | 24 | 5,211 |
| RAL059 | NA | NA | NA | NA | NA | NA | NA | NA |
| RAL91 | CM034800.1 | 647,045 | 652,250 | CM034800.1 | 632,673 | 652,646 | 106 | 5,205 |
| RAL176 | CM034791.1 | 26,085,946 | 26,091,205 | CM034791.1 | 26,038,419 | 26,322,205 | 20 | 5,259 |
| RAL176 | CM034791.1 | 26,085,946 | 26,091,205 | CM034791.1 | 25,408,795 | 26,316,328 | 24 | 5,259 |
| RAL176 | CM034791.1 | 20,918,328 | 20,923,584 | CM034791.1 | 14,159,131 | 24,019,498 | 32 | 5,256 |
| RAL176 | CM034791.1 | 14,905,452 | 14,910,683 | CM034791.1 | 14,159,131 | 24,019,498 | 32 | 5,231 |
| RAL176 | CM034791.1 | 26,093,914 | 26,099,145 | CM034791.1 | 26,038,419 | 26,322,205 | 20 | 5,231 |
| RAL176 | CM034791.1 | 26,093,914 | 26,099,145 | CM034791.1 | 25,408,795 | 26,316,328 | 24 | 5,231 |
| RAL176 | CM034791.1 | 14,340,524 | 14,345,750 | CM034791.1 | 14,159,131 | 24,019,498 | 32 | 5,226 |
| RAL176 | CM034791.1 | 28,810,297 | 28,815,513 | CM034791.1 | 28,771,759 | 29,048,566 | 41 | 5,216 |
| RAL176 | CM034791.1 | 22,320,131 | 22,325,346 | CM034791.1 | 14,159,131 | 24,019,498 | 32 | 5,215 |
| RAL176 | CM034791.1 | 16,010,334 | 16,015,541 | CM034791.1 | 14,159,131 | 24,019,498 | 32 | 5,207 |
| RAL176 | CM034791.1 | 14,919,924 | 14,925,117 | CM034791.1 | 14,159,131 | 24,019,498 | 32 | 5,193 |
| RAL176 | CM034791.1 | 14,893,856 | 14,899,042 | CM034791.1 | 14,159,131 | 24,019,498 | 32 | 5,186 |
| RAL176 | CM034791.1 | 16,015,729 | 16,020,807 | CM034791.1 | 14,159,131 | 24,019,498 | 32 | 5,078 |
| RAL176 | CM034791.1 | 14,934,911 | 14,938,976 | CM034791.1 | 14,159,131 | 24,019,498 | 32 | 4,065 |
| RAL177 | NA | NA | NA | NA | NA | NA | NA | NA |
| RAL375 | NA | NA | NA | NA | NA | NA | NA | NA |
| RAL737 | CM034765.1 | 1,080,088 | 1,085,289 | CM034765.1 | 1,049,206 | 1,620,863 | 78 | 5,201 |
| RAL855 | NA | NA | NA | NA | NA | NA | NA | NA |
| RALISLA001 | CM034749.1 | 26,046,223 | 26,051,454 | CM034749.1 | 26,024,266 | 26,056,883 | 130 | 5,231 |
| RALISLA001 | CM034749.1 | 26,046,223 | 26,051,454 | CM034749.1 | 25,523,728 | 26,335,786 | 24 | 5,231 |
| RALISLA001 | CM034746.1 | 23,293,424 | 23,298,639 | CM034746.1 | 22,047,186 | 23,345,939 | 8_9 | 5,215 |
| STO022 | CM034727.1 | 3,914,775 | 3,920,026 | CM034727.1 | 3,884,495 | 3,975,946 | 36 | 5,251 |
| TOM007 | NA | NA | NA | NA | NA | NA | NA | NA |
| TOM008 | CM034742.1 | 26,304,509 | 26,309,727 | CM034742.1 | 25,253,525 | 27,729,080 | 70 | 5,218 |
| TOM008 | CM034742.1 | 26,937,996 | 26,943,212 | CM034742.1 | 25,253,525 | 27,729,080 | 70 | 5,216 |
| TOM008 | CM034742.1 | 24,403,890 | 24,408,706 | CM034742.1 | 24,278,426 | 24,434,565 | 40 | 4,816 |
| TOM008 | CM034741.1 | 6,489,636 | 6,493,970 | CM034741.1 | 6,468,033 | 6,724,939 | 1 | 4,334 |

Table 7: Overview of the long-read assemblies of diverse insect species analysed in this work.

| order | family | genus | taxon | accession |
|---|---|---|---|---|
| Coleoptera | Coccinellidae | Cryptolaemus | Cryptolaemus.montrouzieri | GCA_013387265.1 |
| Coleoptera | Coccinellidae | Harmonia | Harmonia.axyridis | GCA_011033045.1 |
| Coleoptera | Coccinellidae | Propylea | Propylea.japonica | GCA_013421045.1 |
| Coleoptera | Curculionidae | Listronotus | Listronotus.bonariensis | GCA_014170235.1 |
| Coleoptera | Curculionidae | Sitophilus | Sitophilus.oryzae | GCA_002938485.2 |
| Coleoptera | Elateridae | Limonius | Limonius.californicus | GCA_014611495.1 |
| Coleoptera | Lampyridae | Abscondita | Abscondita.terminalis | GCA_013368085.1 |
| Coleoptera | Lampyridae | Lamprigera | Lamprigera.yunnana | GCA_013368075.1 |
| Coleoptera | Lampyridae | Photinus | Photinus.pyralis | GCA_008802855.1 |
| Coleoptera | Nitidulidae | Aethina | Aethina.tumida | GCA_001937115.1 |
| Coleoptera | Scarabaeidae | Protaetia | Protaetia.brevitarsis | GCA_004143645.1 |
| Coleoptera | Scarabaeidae | Trypoxylus | Trypoxylus.dichotomus | GCA_014905495.1 |
| Coleoptera | Silphidae | Nicrophorus | Nicrophorus.vespilloides | GCA_001412225.1 |
| Collembola | Entomobryidae | Sinella | Sinella.curviseta | GCA_004115045.2 |
| Collembola | Isotomidae | Folsomia | Folsomia.candida | GCA_002217175.1 |
| Collembola | Orchesellidae | Orchesella | Orchesella.cincta | GCA_001718145.1 |
| Diptera | Calliphoridae | Cochliomyia | Cochliomyia.hominivorax | GCA_004302925.1 |
| Diptera | Calliphoridae | Phormia | Phormia.regina | GCA_001735545.1 |
| Diptera | Chironomidae | Belgica | Belgica.antarctica | GCA_000775305.1 |
| Diptera | Culicidae | Aedes | Aedes.albopictus | GCA_001876365.2 |
| Diptera | Culicidae | Aedes | Aedes.aegypti | GCA_002204515.1 |
| Diptera | Culicidae | Anopheles | Anopheles.albimanus | GCA_013758885.1 |
| Diptera | Culicidae | Anopheles | Anopheles.gambiae | GCA_001542645.1 |
| Diptera | Culicidae | Anopheles | Anopheles.funestus | GCA_003951495.1 |
| Diptera | Culicidae | Anopheles | Anopheles.stephensi | GCA_013141755.1 |
| Diptera | Culicidae | Anopheles | Anopheles.coluzzii | GCA_004136515.2 |
| Diptera | Diopsidae | Teleopsis | Teleopsis.dalmanni | GCA_002237135.1 |
| Diptera | Muscidae | Haematobia | Haematobia.irritans | GCA_003123925.1 |
| Diptera | Muscidae | Musca | Musca.domestica | GCA_014843735.1 |
| Diptera | Sarcophagidae | Sarcophaga | Sarcophaga.peregrina | GCA_014635995.1 |
| Diptera | Sciaridae | Bradysia | Bradysia.coprophila | GCA_014529535.1 |
| Diptera | Tephritidae | Bactrocera | Bactrocera.oleae | GCA_001188975.4 |
| Hemiptera | Aleyrodidae | Bemisia | Bemisia.tabaci | GCA_001854935.1 |
| Hemiptera | Aleyrodidae | Trialeurodes | Trialeurodes.vaporariorum | GCA_011764245.1 |
| Hemiptera | Anthocoridae | Orius | Orius.insidiosus | GCA_014119065.1 |
| Hemiptera | Aphididae | Aphis | Aphis.glycines | GCA_009928515.1 |
| Hemiptera | Aphididae | Rhopalosiphum | Rhopalosiphum.maidis | GCA_003676215.3 |
| Hemiptera | Aphididae | Sitobion | Sitobion.miscanthi | GCA_008086715.1 |
| Hemiptera | Delphacidae | Laodelphax | Laodelphax.striatellus | GCA_003335185.2 |
| Hemiptera | Delphacidae | Nilaparvata | Nilaparvata.lugens | GCA_014356525.1 |
| Hemiptera | Liviidae | Diaphorina | Diaphorina.citri | GCA_000475195.1 |
| Hemiptera | Miridae | Apolygus | Apolygus.lucorum | GCA_009739505.2 |
| Hemiptera | Pentatomidae | Euschistus | Euschistus.heros | GCA_003667255.1 |
| Hemiptera | Pseudococcidae | Maconellicoccus | Maconellicoccus.hirsutus | GCA_003261595.1 |
| Hemiptera | Pseudococcidae | Phenacoccus | Phenacoccus.solenopsis | GCA_009761765.1 |
| Hemiptera | Reduviidae | Triatoma | Triatoma.infestans | GCA_011037195.1 |
| Hymenoptera | Apidae | Apis | Apis.mellifera | GCA_003254395.2 |
| Hymenoptera | Apidae | Apis | Apis.dorsata | GCA_009792835.1 |
| Hymenoptera | Apidae | Apis | Apis.mellifera | GCA_003314205.1 |
| Hymenoptera | Apidae | Apis | Apis.cerana | GCA_011100585.1 |
| Hymenoptera | Apidae | Apis | Apis.mellifera | GCA_013841205.1 |
| Hymenoptera | Apidae | Apis | Apis.mellifera | GCA_013841245.1 |
| Hymenoptera | Apidae | Bombus | Bombus.vosnesenskii | GCA_011952255.1 |
| Hymenoptera | Apidae | Bombus | Bombus.bifarius | GCA_011952205.1 |
| Hymenoptera | Apidae | Bombus | Bombus.vancouverensis | GCA_011952275.1 |
| Hymenoptera | Braconidae | Aphidius | Aphidius.gifuensis | GCA_014905175.1 |
| Hymenoptera | Braconidae | Aphidius | Aphidius.ervi | GCA_011426455.1 |
| Hymenoptera | Braconidae | Chelonus | Chelonus.insularis | GCA_013357705.1 |
| Hymenoptera | Braconidae | Lysiphlebus | Lysiphlebus.fabarum | GCA_011426435.1 |
| Hymenoptera | Colletidae | Colletes | Colletes.gigas | GCA_013123115.1 |
| Hymenoptera | Figitidae | Leptopilina | Leptopilina.clavipes | GCA_001855655.1 |
| Hymenoptera | Figitidae | Leptopilina | Leptopilina.boulardi | GCA_011634795.1 |
| Hymenoptera | Formicidae | Camponotus | Camponotus.floridanus | GCA_003227725.1 |
| Hymenoptera | Formicidae | Formica | Formica.selysi | GCA_009859135.1 |
| Hymenoptera | Formicidae | Harpegnathos | Harpegnathos.saltator | GCA_003227715.1 |
| Hymenoptera | Formicidae | Monomorium | Monomorium.pharaonis | GCA_013373865.2 |
| Hymenoptera | Formicidae | Nylanderia | Nylanderia.fulva | GCA_005281655.1 |
| Hymenoptera | Formicidae | Ooceraea | Ooceraea.biroi | GCA_003672135.1 |
| Hymenoptera | Formicidae | Solenopsis | Solenopsis.invicta | GCA_010367695.1 |
| Hymenoptera | Megachilidae | Osmia | Osmia.lignaria | GCA_012274295.1 |
| Hymenoptera | Pteromalidae | Nasonia | Nasonia.vitripennis | GCA_009193385.2 |
| Hymenoptera | Pteromalidae | Pteromalus | Pteromalus.puparum | GCA_012977825.2 |
| Hymenoptera | Vespidae | Polistes | Polistes.metricus | GCA_010416925.1 |
| Hymenoptera | Vespidae | Polistes | Polistes.fuscatus | GCA_010416935.1 |
| Hymenoptera | Vespidae | Vespa | Vespa.mandarinia | GCA_014083535.1 |
| Lepidoptera | Carposinidae | Carposina | Carposina.sasakii | GCA_014607495.2 |
| Lepidoptera | Crambidae | Chilo | Chilo.suppressalis | GCA_004000445.1 |
| Lepidoptera | Crambidae | Cnaphalocrocis | Cnaphalocrocis.medinalis | GCA_014851415.1 |
| Lepidoptera | Hesperiidae | Epargyreus | Epargyreus.clarus | GCA_014595695.1 |
| Lepidoptera | Lasiocampidae | Dendrolimus | Dendrolimus.punctatus | GCA_012273795.1 |
| Lepidoptera | Noctuidae | Heliothis | Heliothis.virescens | GCA_002382865.1 |
| Lepidoptera | Noctuidae | Spodoptera | Spodoptera.exigua | GCA_011316535.1 |
| Lepidoptera | Noctuidae | Spodoptera | Spodoptera.frugiperda | GCA_011064685.1 |
| Lepidoptera | Noctuidae | Trichoplusia | Trichoplusia.ni | GCA_003590095.1 |
| Lepidoptera | Nymphalidae | Maniola | Maniola.jurtina | GCA_009667785.1 |
| Lepidoptera | Pieridae | Colias | Colias.croceus | GCA_009982905.1 |
| Lepidoptera | Psychidae | Eumeta | Eumeta.japonica | GCA_005406025.1 |
| Lepidoptera | Pyralidae | Galleria | Galleria.mellonella | GCA_004355975.1 |
| Lepidoptera | Saturniidae | Antheraea | Antheraea.mylitta | GCA_014332785.1 |
| Lepidoptera | Saturniidae | Samia | Samia.ricini | GCA_014132275.1 |
| Lepidoptera | Sphingidae | Hyles | Hyles.vespertilio | GCA_009982885.1 |
| Lepidoptera | Sphingidae | Manduca | Manduca.sexta | GCA_014839805.1 |
| Lepidoptera | Tortricidae | Cydia | Cydia.pomonella | GCA_003425675.2 |
| Orthoptera | Gryllidae | Teleogryllus | Teleogryllus.occipitalis | GCA_011170035.1 |
| Siphonaptera | Pulicidae | Ctenocephalides | Ctenocephalides.felis | GCA_003426905.1 |
| Thysanoptera | Thripidae | Thrips | Thrips.palmi | GCA_012932325.1 |
| Trichoptera | Hydropsychidae | Hydropsyche | Hydropsyche.tenuis | GCA_009617725.1 |
| Trichoptera | Polycentropodidae | Plectrocnemia | Plectrocnemia.conspersa | GCA_009617715.1 |
| Trichoptera | Stenopsychidae | Stenopsyche | Stenopsyche.tienmushanensis | GCA_008973525.1 |

### References

S. F. Altschul, W. Gish, W. Miller, E. W. Myers, and D. J. Lipman. Basic local alignment search tool. *Journal of Molecular Biology*, 215(3):403–410, 1990.

M. Chakraborty, J. J. Emerson, S. J. Macdonald, and A. D. Long. Structural variants exhibit widespread allelic heterogeneity and shape variation in complex traits. *Nature Communications*, 10(1):4872, 2019.

M. Chakraborty, C. Chang, D. Khost, J. A. J Vedanayagam, Y. Liao, K. Montooth, C. Meiklejohn, A. Larracuente, and J. Emerson. Evolution of genome structure in the *Drosophila simulans* species complex. *Genome Research*, 31:380–396, 2021.

C. E. Ellison and W. Cao. Nanopore sequencing and Hi-C scaffolding provide insight into the evolutionary dynamics of transposable elements and piRNA production in wild strains of *Drosophila melanogaster*. *Nucleic Acids Research*, 48(1):1–14, 2020.

L. S. Gramates, J. Agapite, H. Attrill, B. R. Calvi, M. A. Crosby, G. Dos Santos, J. L. Goodman, D. Goutte-Gattat, V. K. Jenkins, T. Kaufman, et al. Flybase: a guided tour of highlighted features. *Genetics*, 220 (4):iyac035, 2022.

J. K. Grenier, J. R. Arguello, M. C. Moreira, S. Gottipati, J. Mohammed, S. R. Hackett, R. Boughton, A. J. Greenberg, and A. G. Clark. Global diversity lines–a five-continent reference panel of sequenced *Drosophila melanogaster* strains. *G3: Genes, Genomes, Genetics*, 5(4):593–603, 2015.

R. A. Hoskins, J. W. Carlson, K. H. Wan, S. Park, I. Mendez, S. E. Galle, B. W. Booth, B. D. Pfeiffer, R. A. George, R. Svirskas, et al. The Release 6 reference sequence of the *Drosophila melanogaster* genome. *Genome Research*, 25(3):445–458, 2015.

E. G. King, C. M. Merkes, C. L. McNeil, S. R. Hoofer, S. Sen, K. W. Broman, A. D. Long, and S. J. Macdonald. Genetic dissection of a model complex trait using the Drosophila Synthetic Population Resource. *Genome Research*, 22(8):1558–1566, 2012.

J. D. Lange, H. Bastide, J. B. Lack, and J. E. Pool. A Population Genomic Assessment of Three Decades of Evolution in a Natural Drosophila Population. *Molecular Biology and Evolution*, 39(2), 2021.

Q. Long, F. A. Rabanal, D. Meng, C. D. Huber, A. Farlow, A. Platzer, Q. Zhang, B. J. Vilhjálmsson, A. Korte, V. Nizhynska, et al. Massive genomic variation and strong selection in *Arabidopsis thaliana* lines from Sweden. *Nature genetics*, 45(8):884–890, 2013.

T. F. Mackay, S. Richards, E. A. Stone, A. Barbadilla, J. F. Ayroles, D. Zhu, S. Casillas, Y. Han, M. M. Magwire, J. M. Cridland, et al. The *Drosophila melanogaster* genetic reference panel. *Nature*, 482(7384): 173–178, 2012.

H. Quesneville, C. M. Bergman, O. Andrieu, D. Autard, D. Nouaud, M. Ashburner, and D. Anxolabéhère. Combined evidence annotation of transposable elements in genome sequences. *PLoS Computational Biology*, 1(2):166–175, 2005.

G. E. Rech, S. Radío, S. Guirao-Rico, L. Aguilera, V. Horvath, L. Green, H. Lindstadt, V. Jamilloux, H. Quesneville, and J. González. Population-scale long-read sequencing uncovers transposable elements associated with gene expression variation and adaptive signatures in drosophila. *Nature Communications*, 13(1):1948, 2022.

F. Schwarz, F. Wierzbicki, K.-A. Senti, and R. Kofler. Tirant Stealthily Invaded Natural *Drosophila melanogaster* Populations during the Last Century . *Molecular Biology and Evolution*, 38(4):1482–1497, 2021.

A. Shumate and S. Salzberg. Liftoff: accurate mapping of gene annotations. *Bioinformatics*, 37(12):1639–1643, 2021.

A. F. A. Smit, R. Hubley, and P. Green. RepeatMasker Open-3.0, 1996-2010. URL `http://www.repeatmasker.org`.

S. Srivastav, C. Feschotte, and A. G. Clark. Rapid evolution of piRNA clusters in the *Drosophila melanogaster* ovary. *bioRxiv*, 2023.

F. Wierzbicki, F. Schwarz, O. Cannalonga, and R. Kofler. Novel quality metrics allow identifying and generating high-quality assemblies of piRNA clusters. *Molecular Ecology Resources*, 2021. doi: 10.1111/1755-0998.13455.
